# A dynamic role for the apical extracellular matrix in cell migration during *Drosophila* abdominal morphogenesis

**DOI:** 10.64898/2026.11.18.767616

**Authors:** Jianyi Mai, Philip Wijesinghe, Jochen Kursawe, Marcus Bischoff

**Affiliations:** Biomolecular Sciences Research Complex, University of St Andrews, North Haugh, St Andrews, KY16 9ST, United Kingdom; Centre of Biophotonics, University of St Andrews, North Haugh, St Andrews, KY16 9SS, United Kingdom; SUPA, School of Physics and Astronomy, University of St Andrews, St Andrews, KY16 9SS, United Kingdom; School of Mathematics, University of St Andrews, St Andrews, KY16 9SS, United Kingdom

## Abstract

Cell migration plays a central role in development, ensuring cells reach their correct destinations. Many migratory cell types require interaction with the extracellular matrix (ECM) to generate traction. While the basal ECM (bECM) is well known for its role in cell motility, the apical ECM (aECM) has mostly been shown to provide structural and mechanical support during morphogenesis and remains less understood. In *Drosophila*, the aECM is primarily composed of the giant protein Dumpy (Dpy), which interacts with the Zona Pellucida (ZP) domain protein Piopio (Pio). In this study, we elucidate the role of the aECM in the formation of the adult abdominal epidermis of *Drosophila*. During this process, the larval epithelial cells (LECs) transition from a stationary to a mobile state, in which they migrate directionally, before apically constricting and undergoing apoptosis. Using *in vivo* 4D microscopy, we examined the role of the aECM in the dynamic behaviour of the LECs. Initially, a layer of Dpy and Pio is located apically to the stationary LECs. As migration begins, the LECs leave behind this early layer while depositing an additional, morphologically distinct layer of Dpy. Knock-down of Dpy or Pio leads to the detachment of stationary LECs from the early layer and, subsequently, to impaired LEC migration. Our results reveal that the aECM transitions from acting as an anchoring scaffold to supporting cell migration. This functional change is closely linked to the change in LEC behaviour. Together, these findings identify the aECM as a substrate for cell migration and highlight a dynamic role of the aECM during morphogenesis.

## Introduction

Cell migration plays a central role in development, actively shaping tissues and organs by repositioning cells to new locations (1). This often involves cells transitioning from a stationary to a mobile state, enabling dynamic tissue remodelling, as exemplified by the migration of *Drosophila* border cells (2) and primordial germ cells (3), as well as vertebrate neural crest cells (4) and gastrulation movements (1). Migration of many cell types requires cell-substrate adhesion to generate traction, primarily mediated by integrins interacting with the basal extracellular matrix (bECM) (5). The bECM not only provides structural support but also transmits mechanical cues and guidance signals that direct collective and individual cell movements across various morphogenetic processes (6,7).

Besides the bECM, cells also produce an apical ECM (aECM) (8,9). While bECM is primarily located at internal faces of cells and tissues, aECM generally faces the external environment, forming the outermost skin layer and lining the surfaces of internal tubular epithelia and endothelia (8). For example, aECM constitutes the cuticle of *C. elegans* (10) and *Drosophila* (11). Other aECMs can be found in the excretory canal and vulva of *C. elegans* (12). aECMs are also found in vertebrates, where they form the vascular glycocalyx, line the gastrointestinal tract and upper airways, and constitute the alveolar surfactant in the lungs (8). Furthermore, aECM is found in vertebrate inner ear, where it is involved in sensory transduction (13).

While the bECM has been extensively studied, the aECM remains less well understood. In *Drosophila*, the main component of the aECM is Dumpy (Dpy), a very large 2.5 MDa protein, which comprises 308 epidermal growth factor (EGF) modules interspersed with 185 unique DPY repeats, followed by a conserved Zona Pellucida (ZP) domain, a proximal transmembrane (TM) region, and a short cytoplasmic tail (14). The ZP domain is thought to mediate the polymerisation of ZP domain proteins into matrices and filaments (15). A consensus furin cleavage site between the ZP and TM domains allows proteolytic shedding of the Dpy ectodomain (16). Dpy plays a role in several morphogenetic processes, e.g. promoting tube expansion during tubular morphogenesis of the trachea (17,18) and salivary glands (19). Dpy is involved in shaping the wings, legs and antennae by linking epithelial cells to the cuticle, with remodelling of the aECM being crucial for driving morphogenesis (20–22). Dpy fibres provide resistance at the points where epithelial cells adhere to the cuticle (14). Additionally, under mechanical tension, Dpy forms filaments that tether tendon cells, which connect the flight muscles to the cuticle (16).

Dpy interacts with the ZP-domain protein Piopio (Pio) (15,17). The interaction between Dpy and Pio likely depends on their ZP-domains (15,17,23). Pio is involved in Dpy secretion (18,19). Similar to Dpy, Pio is essential for both tracheal network formation (17,18,24) and salivary gland development (19). Furthermore, Pio contributes to cell-aECM adhesion in *Drosophila* wings (25) and mediates connections between tendon cells and the chitinous exoskeleton in embryos and larvae (26). Additionally, Dpy and Pio are part of an external scaffold that is crucial during *Drosophila* corneal lens morphogenesis (27).

Overall, much of what is known about aECM highlights its role in providing structural and mechanical support during morphogenesis. The secreted aECM in tubular lumen serves as an elastic scaffold, providing tensile stress to stabilise the growing trachea (18,24). Attachment of the cells to the aECM holds the tensile stresses under high mechanical forces during tissue and organ deformation (16,27,28), wing expansion (20–22), and locomotion (26).

In this study, we investigated the role of the aECM in the morphogenesis of the adult abdominal epidermis in *Drosophila*. Here, diploid histoblasts replace polyploid larval epithelial cells (LECs) to form the adult epidermis (29–32). Initially stationary, the LECs become mobile und undergo directed migration, before they apically constrict and undergo apoptosis (Fig. 1A) (33). From the onset of abdominal morphogenesis, the bECM beneath the LECs and histoblasts undergoes degradation (34,35), leaving only the aECM. While migrating, LECs form apical lamellipodia, suggesting that they crawl on an apical substrate, possibly the aECM (33). However, the role of the aECM in abdominal morphogenesis remains unclear.

**Figure 1.**
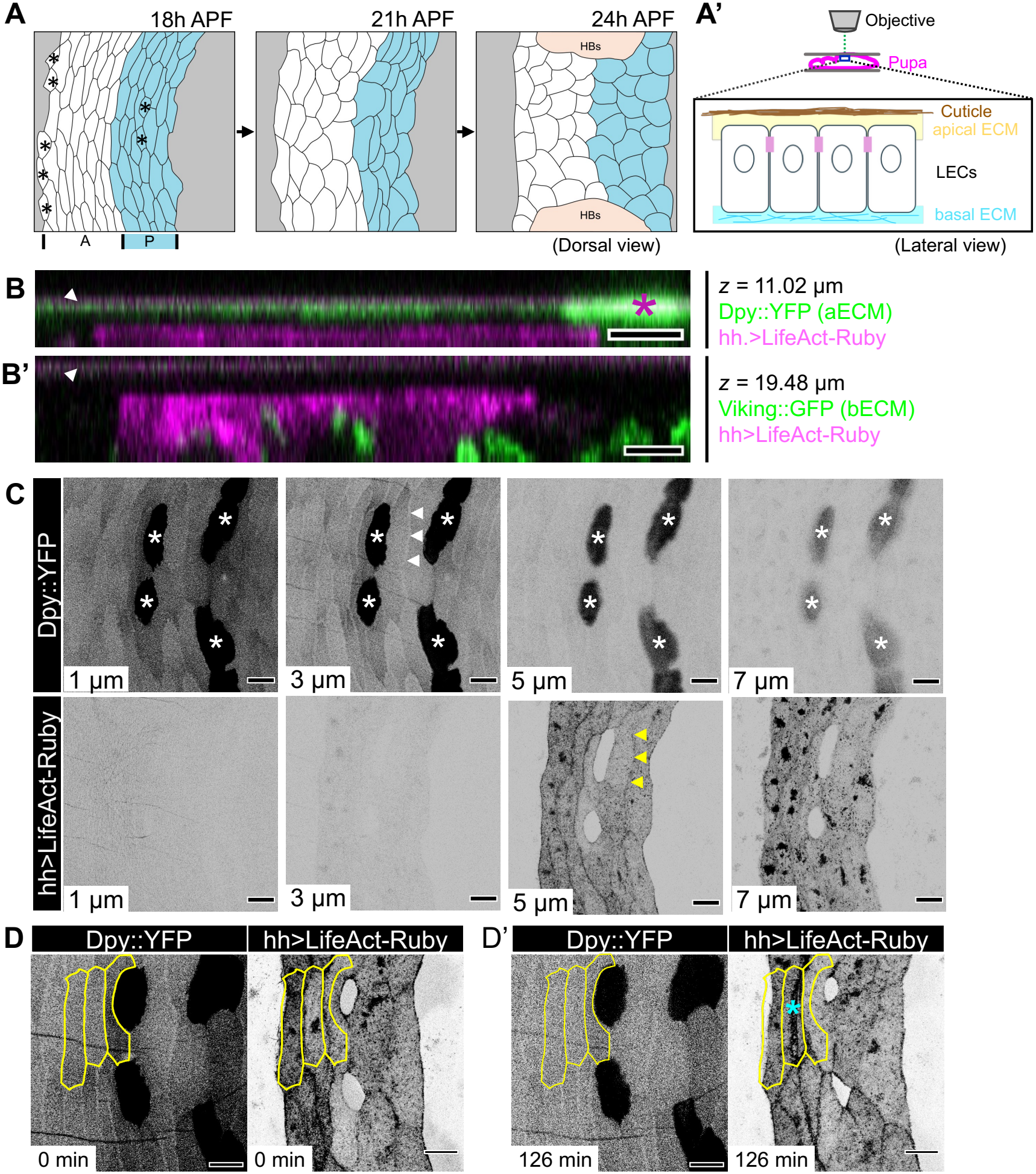
Dpy::YFP localisation in early stationary LECs. The early layer of Dumpy is cleaved and remains in the aECM as LECs begin early apoptosis and posterior migration. **(A)** Scheme of morphogenesis of the adult abdominal epidermis. LECs transition from stationary to migratory behaviour. They form crescent-shaped protrusions, migrate, constrict, and undergo apoptosis, while they are replaced by histoblasts (HBs). A (white) and P (blue) compartments are shown; asterisks mark tendon cells. **(A’)** *In vivo* 4D imaging of the dorsal side of the abdominal epidermis. Lateral view of LECs and the ECM. **(B, B’)** Orthogonal views of aECM and bECM of LECs. *XZ* sections are average projections of five slices. LifeAct-Ruby labels P compartment LECs. **(B)** Dpy::YFP localises to the aECM, near the pupal cuticle, which is visible due to its autofluorescence in the magenta channel (white arrowhead) (*N*=10 pupae). Dpy::YFP is particularly enriched apically above tendon cells (asterisk). **(B’)** Viking::GFP localises to the basal side of the tissue (*N*=6 pupae). **(C)** *Z*-stack images of LECs expressing Dpy::YFP homozygously. LifeAct-Ruby labels the P compartment. Dpy::YFP localises close to the cuticle (at 1 μm), the aECM (at 3 μm), the apical membrane (at 5 μm) and the cytoplasm (at 7 μm). Dpy is enriched in the tendon cell aECM (white asterisks) and absent above adherens junctions (white arrowheads; yellow arrowheads highlight the corresponding junction) (*N*=7 field of views (FOVs) in 6 pupae). **(D, D’)** The early Dpy::YFP layer is cleaved and left behind in the aECM as LECs begin posterior migration and undergo apoptosis (*N*=9 pupae). Time-lapse images of LECs expressing Dpy::YFP heterozygously. LifeAct-Ruby labels P compartment LECs. **(D)** LECs at the onset of migration. The initial Dpy::YFP layer reflects the original LEC cell shape, with Dpy::YFP being absent above the adherens junctions (three cells outlined in yellow). LifeAct-Ruby shows that cells have begun changing shape. **(D’)** 126 min later, the pattern of the early Dpy::YFP layer remains fixed in aECM (yellow lines) while cells change shape below, with the middle of the marked cells strongly reducing apical area while undergoing apoptosis (cyan asterisks). In all images, anterior is to the left. Asterisks indicate tendon cells. Scale bars, 10 μm (B, B’); 20 μm (C, D).

Here, we identified an active role of the aECM in abdominal morphogenesis. During the early stages of morphogenesis, while the LECs remain stationary, Dpy and Pio are present in the aECM and are localised to the apical side of the LECs. Once LECs started migrating, LECs secrete an additional layer of Dpy and Pio with which they interact with during their migration. Both Dpy and Pio knock-down result in early LEC detachment and impaired LEC migration. Our data reveal that the aECM assumes distinct functions during the early and late stages of abdominal morphogenesis. Initially, it anchors stationary LECs to the cuticle, while later it serves as a substrate for their migration. This transition is associated with the dynamic remodelling of the aECM that is driven by ongoing deposition of Dpy carried out by the LECs. This highlights a role for the aECM as a substrate for LEC migration.

## Results

### Dpy is part of the aECM of LECs

To study the role of the aECM in abdominal morphogenesis, we examined the main component of the *Drosophila* aECM, Dpy, using a Dpy::YFP protein trap, which tags Dpy in its extracellular domain (20,36). Dpy::YFP localised apically to the LECs (Fig. 1A’, B), while Collagen IV (Viking::GFP) (37), a bECM component, was localised basally (Fig. 1B’). *Z*-stacks from before the onset of morphogenesis showed Dpy::YFP in the aECM near the cuticle, the apical LEC membrane and in the cytoplasm (Fig. 1C). A large amount of Dpy::YFP localised to the tendon cells and the overlying aECM (Fig. 1C), highlighting the role of Dpy in muscle attachment also in the abdominal epidermis. While the localisation of Dpy::YFP was uniform close to the cuticle, it was absent in the apical cell area close to the cell junctions (Fig. 1C; Suppl. Movie S1), indicating that Dpy localisation is restricted to the apical plasma membrane. The Dpy::YFP observed at this stage we refer to as the ‘early’ layer.

### LECs leave behind the early layer of Dpy when they begin to migrate

During abdominal morphogenesis, some LECs reduce apical area, extrude from the epithelium and undergo apoptosis, while the remaining cells transit from stationary to migratory behaviour (33). We found that the pattern of Dpy::YFP in the aECM did not change when LECs begun to extrude and move (Fig. 1D). Instead, the early layer of extracellular Dpy::YFP was left behind, appearing like a footprint (Fig. 1D). This is likely due to Dpy::YFP being cleaved, leaving the extracellular domain including the YFP-tag behind. This suggests that early LECs initially attach to the aECM and this attachment is severed at the start of morphogenesis.

### LECs secrete a new layer of Dpy during their migration

During the imaging of LEC migration, the fluorescence of the early Dpy::YFP layer decreased over time, due to photobleaching (Fig. 2Aii, B). However, the fluorescence intensity at the interface between the aECM and the LECs increased (Fig. 2Aiii, B; Suppl. Movie S1). This suggests that migrating LECs secrete a new layer of Dpy::YFP, which is deposited into the aECM below the early layer. We refer to this layer as the ‘late’ layer. The late layer is qualitatively different from the early layer: the early Dpy::YFP layer displayed a punctate pattern (Fig. 2C), whereas the late layer exhibited a fibrous appearance (Fig. 2C’), which became more complex over time (Suppl. Movie S1).

**Figure 2.**
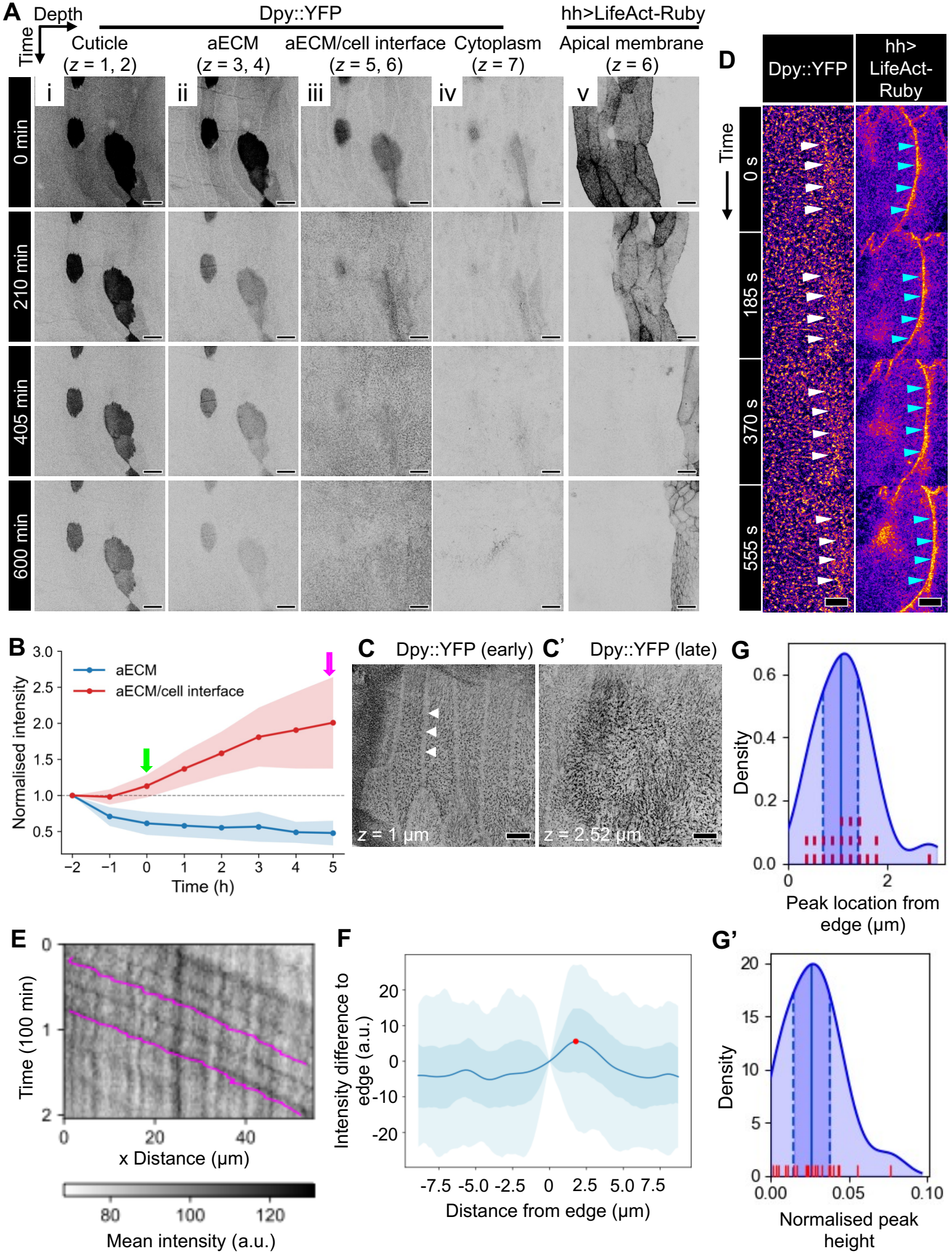
A new, late layer of Dpy::YFP is secreted into the aECM during LEC migration. **(A)** Time-lapse of *z*-stacks (rows=time, columns=depth) showing Dpy::YFP dynamics with LifeAct-Ruby marking P compartment LECs (column (v)). LECs exhibit posterior migration (210 min), dorsal migration and apical constriction (405 min), leading to complete LEC replacement by the histoblasts (600 min). A new, late layer of Dpy::YFP forms at the LEC-aECM interface (iii), while the pattern of the early layer (i, ii) and cytoplasmic signal (iv) change only due to photobleaching (*N*=10 pupae). **(B)** Quantification of the normalised fluorescence intensity of Dpy::YFP at the *z* levels corresponding to the aECM (column ii in A) and aECM/cell interface (iii in A), respectively, from two hours before the start of posterior migration (green arrow) until dorsal migration begins (cyan arrow). The aECM signal bleaches over time, whereas the signal at the aECM/cell interface increases (n=7 pupae; shading indicates standard deviation). **(C, C’)** Fixed LECs showing the apical punctate Dpy::YFP pattern of the early layer, which is absent above adherens junctions (white arrowheads, C), and the fibrous late Dpy::YFP layer (C’) (*N*=4 FOVs in 2 pupae). **(D)** A wave-like signal in the late Dpy::YFP layer (white arrowheads) moves with LECs across the tissue (*N*=17 pupae; 185 s/frame). LifeAct-Ruby shows lamellipodium (cyan arrowheads). Fire look-up table (LUT) used. (**E-G**) Analysis of the position of the wave-like Dpy::YFP signal relative to the LECs’ leading edge. **(E)** Kymograph shows the wave-like Dpy::YFP signal travelling anterior-posterior (black diagonal lines; inverted grey scale LUT). The average position of the leading edge is indicated for two cells in pink. It trails the Dpy::YFP wave. **(F, G)** Quantification of the position of the Dpy::YFP wave relative to the lamellipodial edge. **(F)** Representative example of the average intensity profile of the Dpy::YFP signal relative to the leading edge for one lamellipodium. The Dpy::YFP signal has a pronounced peak (red dot) in front of the leading edge (0 on x-axis). Shaded areas present 50% and 90% equal tail intervals. **(G)** Distributions of Dpy::YFP peak location (G) and normalised peak height (G’) (*N* = 21 lamellipodia; median, 25%, 75% quantiles highlighted). Underlying data points represented in red. Mean Dpy::YFP peak location is 1.15 µm in front of the leading edge, and mean peak height is 2.7% above the average Dpy::YFP fluorescence intensity at the leading edge. In all images, anterior is to the left. Scale bars in 20 μm (A); 10 μm (C, D).

### LECs interact with the aECM during their migration

We next asked whether migrating LECs interact with the aECM by visualising Dpy::YFP dynamics *in vivo* using 4D microscopy. We observed crescent-shaped waves of high fluorescence intensity moving across the late Dpy::YFP layer (Fig. 2D, E; Suppl. Movies S1, S2). The Dpy-YFP signal was observed posterior to the lamellipodia of the LECs (Fig. 2E, F), with the waves travelling an average of 1.15 µm ahead of the lamellipodium (Fig. 2G) and exhibiting a mean intensity increase of approximately 3% (Fig. 2G’).

To investigate further whether LECs adhere to the aECM, we assessed whether LECs deform the aECM when they undergo pulsed contractions (38,39). Using DEFORM-Net (40), we quantified the displacement vector fields corresponding to the motion of features marked by the Dpy::YFP and LifeAct-Ruby fluorescence. Deformation of the late Dpy::YFP layer correlated with the periodic assembly and disassembly of actin foci in the pulsing cells (*N*=11 LECs in 7 pupae; Suppl. Fig. S1; Suppl. Movie S3). These results demonstrate that migratory LECs interact with and pull on the late Dpy::YFP layer.

### Knock-down of *dpy* impairs LEC migration

To study the role of the two layers of Dpy in abdominal morphogenesis, we knocked down *dpy* and asked how LEC behaviour was affected. First, to study the defects in LEC behaviour in *dpy-RNAi* pupae, we expressed *dpy-RNAi* only in the posterior (P) compartment LECs (*N*=10 pupae). This resulted in two phenotypic groups, in both of which, the early layer of Dpy::YFP was reduced above the P compartment (Fig. 3A’, A’’). This reduction was not observed in the A compartment (Fig. 3A’, A’’), which did not express the *dpy-RNAi*, suggesting that the RNAi targets *dpy*. The strength of the phenotype corresponded to the length of time the RNAi was expressed before imaging (i.e. the length of incubation at 29ºC; 32-73 h [weaker phenotype]; 73-120 h [stronger phenotype]). (i) In weaker phenotypes *(N*=7/10 pupae), LEC posterior migration still took place (Fig. 3A’), however, cells migrated more slowly compared to control cells (Fig. 3A’’’). As the cells migrated, the wild-type A compartment LECs deposited a new layer of Dpy::YFP into the aECM of the P compartment territory that had reduced Dpy::YFP due to the *dpy-RNAi* (Fig. 3A’; Suppl. Movie S4). This highlights the deposition of the new, ‘late’ layer of Dpy::YFP and shows that the deposition is independent of the presence of an early Dpy::YFP layer. (ii) In stronger *dpy-RNAi* phenotypes (*N*=3/10 pupae), the migratory behaviour of the P compartment LECs was significantly impaired (Fig. 3A’’’; Suppl. Movie S5). Also, LECs showed a narrow cell morphology along the anterior-posterior (a-p) axis at the stationary stage, resulting in a narrow P compartment (Fig. 3A’’; Suppl. Fig. S2). Notably in these cases, both A and P compartment LECs showed little posterior movement, suggesting that the P compartment LECs’ failure to migrate likely restricted the movement of the A compartment cells (Fig. 3A’’; Suppl. Movie S5). As LECs migrated very little, the reduction of Dpy::YFP above the P compartment remained visible throughout morphogenesis, as the A compartment LECs did not advance into this region to deposit the new, ‘late’ layer of Dpy::YFP (Fig. 3A’’). In both weak and strong phenotypes, abdominal closure still took place, as apical constriction, extrusion, and apoptosis created dorsally directed LEC movement, while histoblasts replaced the LECs (Suppl. Movies S4, S5). Thus, the observed phenotype is specific to LEC migration.

**Figure 3.**
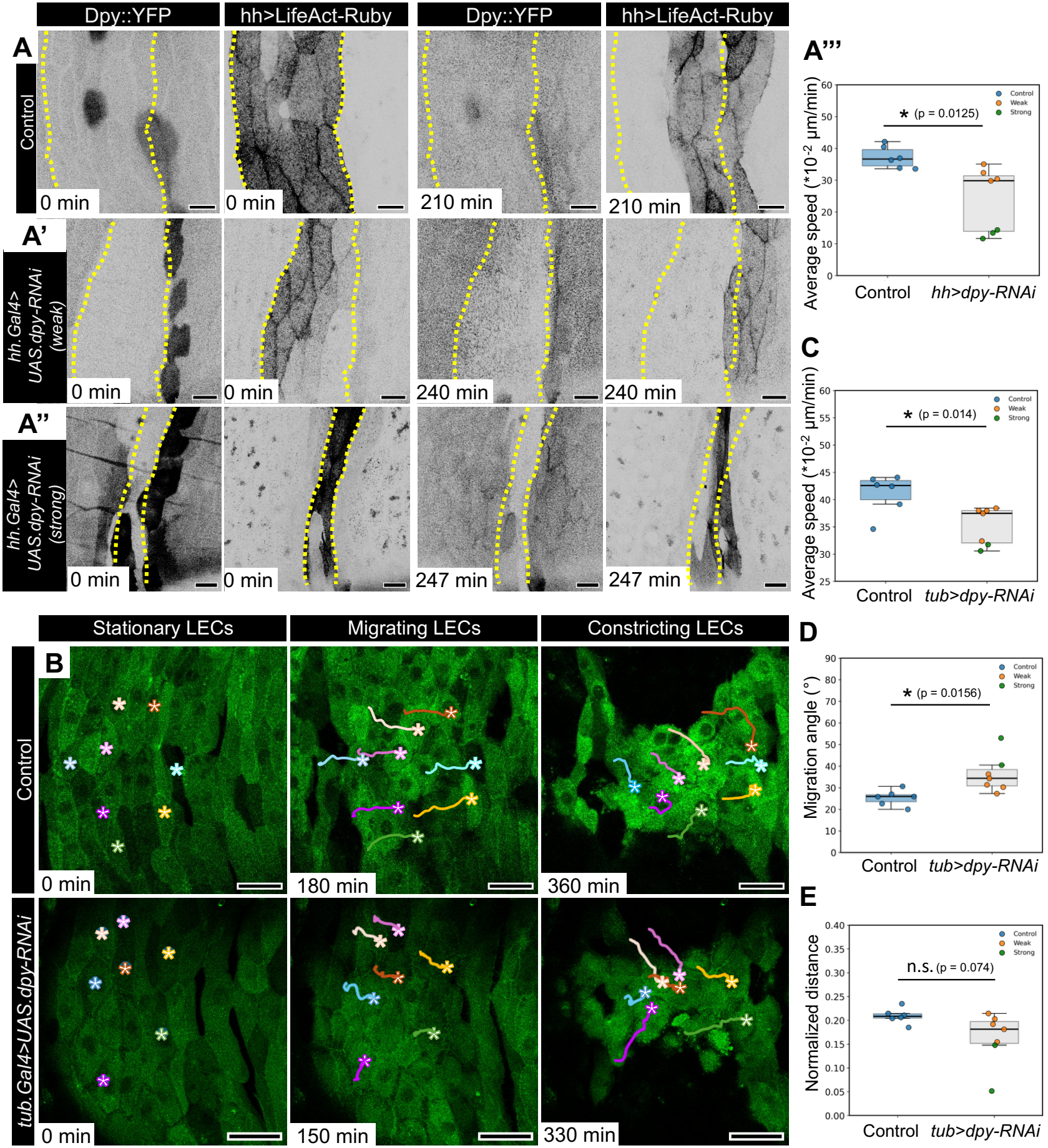
Knock-down of dpy disrupts LEC migration (A-A’’) Knock-down of *dpy* in P compartment LECs. LECs express Dpy::YFP; P compartment cells also express LifeAct-Ruby and *dpy-RNAi*. Yellow dashed lines indicate the location of the P compartment before migration. **(A)** Control shows the early Dpy::YFP layer before migration (0 min) and the late Dpy::YFP layer during migration (210 min) (*N*=10 pupae). **(A’)** Weak *dpy-RNAi* shows reduced the early Dpy::YFP layer above P compartment LECs (0 min). Migrating anterior LECs deposit a late Dpy::YFP layer in the area that has previously been occupied by the P compartment (240 min) (*N*=7/10 pupae). **(A’’)** Strong *dpy-RNAi* shows a reduced early Dpy::YFP layer above P compartment LECs and narrow P compartment LECs (0 min), as well as strongly impaired posterior migration of P compartment LECs, as indicated by the yellow hatched lines (247 min). As LECs are not migrating, the area above the P compartment remains with little Dpy::YFP (*N*=3/10 pupae). **(A’’’)** Posterior migration defects in *dpy-RNAi* evaluated by migration speed. Knock-down of *dpy* significantly reduces speed (*N*=7 pupae) compared to the control (*N=*6 pupae). The strength of the phenotype corresponds to the length of time the RNAi was expressed before imaging (weaker phenotype: 32-50 h, orange data points (*N*=4 pupae); stronger phenotype: 72-120 h, green data points, *N*=3 pupae). **(B-E)** *dpy-RNAi* in all LECs impairs LEC posterior migration. **(B)** Time-lapse images of abdominal morphogenesis showing migration and constriction of LECs. LECs are labelled with mCD8-GFP. Trajectories of selected LECs shown. Compared to control pupae (*N*=10 pupae), LEC posterior migration is impaired in *dpy-RNAi* pupae (*N*=7 pupae). **(C-E)** Using the cell tracking data, average posterior migration speed (**C**), average migration direction **(D)** and average normalised migration distance **(E)** were calculated (see Methods). Compared to the control (*N*=6 pupae), *dpy-RNAi* reduces speed and decreases directionality significantly, but affects normalised distance only in strong phenotypes (*N*=7 pupae). The strength of the phenotype corresponds to the length of time the RNAi was expressed before imaging (weaker phenotype: 27 h, orange data points (*N*=5 pupae); stronger phenotype: 53 h, green data points (*N*=2 pupae). In all images, anterior is to the left. Scale bars, 20 μm (A); 50 μm (B). Box plots show median and interquartile range. Whiskers extend to 1.5x the interquartile range. A’’’, D, E, Welch’s t-test; C, Mann-Whitney U test.

Secondly, we asked whether *dpy-RNAi* expression in all LECs, using a *tub*.*Gal4* FLP-out cassette (31,41), also affected LEC migration. Using manual cell tracking with SIMI Biocell (42), we analysed LEC migratory behaviour throughout morphogenesis, comparing control (Suppl. Fig. S3) with *dpy-RNAi* pupae (Suppl. Fig. S4A). The analysis revealed that, during posterior migration, *dpy-RNAi* LECs migrated at a slower rate and were less directional than control cells (Fig. 3B, C, D). There was no significant difference in the migration distance between control and *dpy-RNAi* cells, however, strong phenotypes showed a reduction of migration distance (Fig. 3E). The strength of the migration phenotype corresponded to the length of time the RNAi was expressed before imaging (i.e. the time elapsed since FLP-out activation; 27 h [weaker phenotype]: *N* = 5/7; 53 h [stronger phenotype]: *N* = 2/7; Fig. 3C, D, E). Similar to the *hh*.*Gal4* experiments, abdominal closure still took place, although LEC migration was impaired (Suppl. Movies S6, S7).

Taken together, our results demonstrate that Dpy is necessary for LEC migration. The observed migration phenotypes are consistent with Dpy acting as a substrate that facilitates the normal migration of LECs.

### Pio is required for LEC migration

To gain further insights into the role of the aECM in LEC migration, we turned to Pio. Pio has previously been implicated in LEC migration, however, the phenotypic description was qualitative (33). We confirmed that *pio* is required in LEC migration, by quantifying the impacts of *pio-RNAi* in all LECs (*tub*.*Gal4*) on posterior migration by manually tracking cell movement (Fig. 4A’; Suppl. Fig. S4B; Suppl. Movie S8). The knock-down of *pio* significantly impaired LEC posterior migration, resulting in a significantly slower migration speed, an abnormal migration direction, and a shorter migration distance (Fig. 4B-D). Similar to the *dpy-RNAi* experiments, LECs still moved dorsally, as apical constriction, extrusion, and apoptosis of LECs decreased the overall LEC tissue area (Suppl. Movie S8). The phenotype of knock-down of *pio* (Fig. 4B-D; Suppl. Fig. S4B) is more severe than the phenotype of the knock-down of *dpy* (Fig. 3C-E; Suppl. Fig. S4A). Overall, our results show that Pio is necessary for LEC posterior migration, and the similarities in phenotype between Dpy and Pio strongly suggest they are part of the same mechanism.

**Figure 4.**
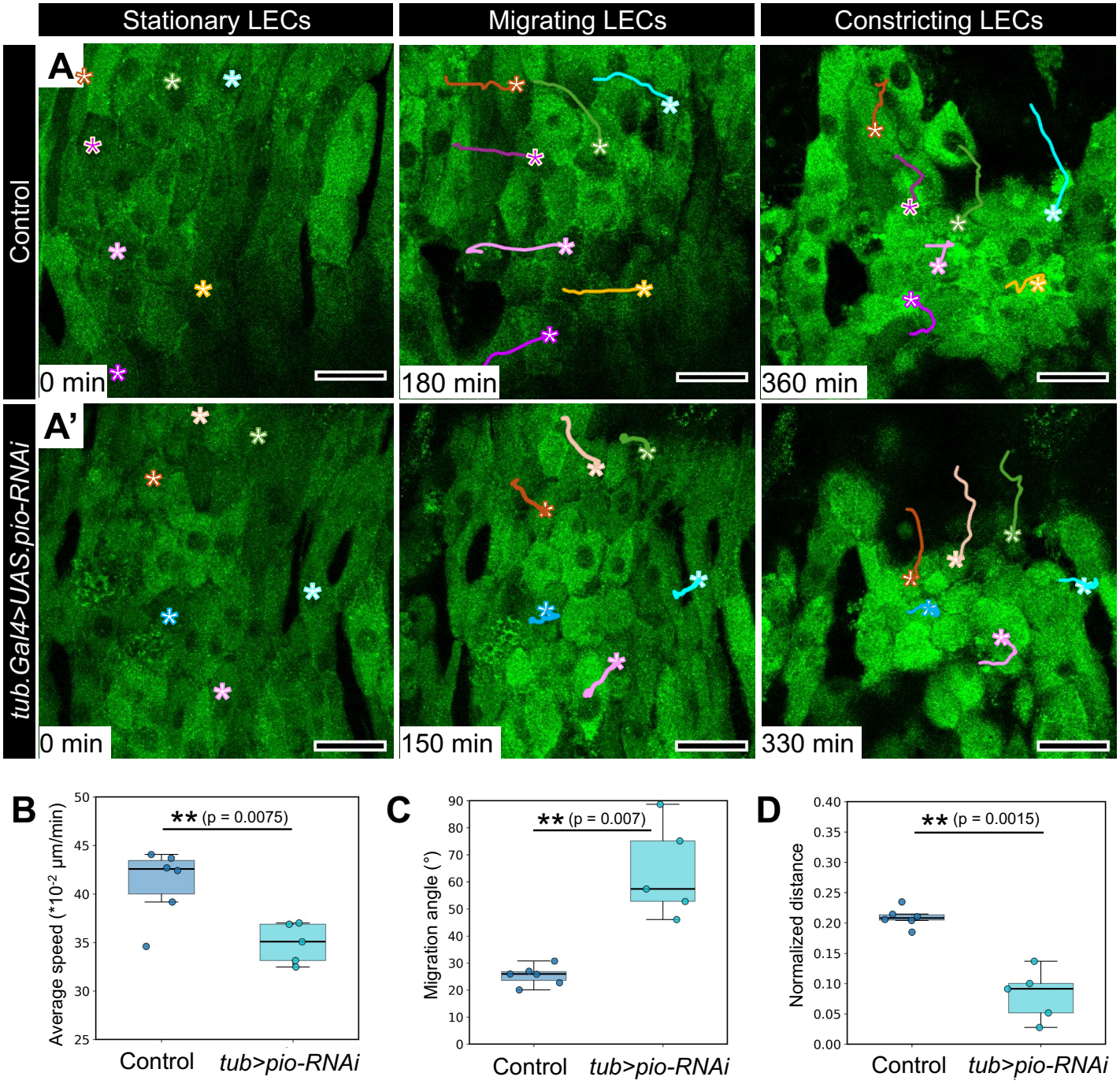
Knock-down of *pio* interferes with LEC posterior migration. Expressed in all LECs, *pio-RNAi* impairs LEC posterior migration. **(A, A’)** Time-lapse images of abdominal morphogenesis showing migration and constriction of LECs. LECs are labelled with mCD8-GFP. Trajectories of selected LECs shown. Compared to control pupae (A; *N*=10 pupae), LEC posterior migration is impaired in *pio-RNAi* pupae (A’; *N*=13 pupae). **(B-D)** Using the cell tracking data, average posterior migration speed **(B)**, average migration direction **(C)** and average normalised migration distance **(D)** were calculated (see Methods). Control data presented in Fig. 3C-E (*N*=6 pupae), shown again here for convenience. Compared to the control, *pio-RNAi* reduces speed and normalised distance, and decreases directionality (*N*=5 pupae). In all images, anterior is to the left. Scale bars, 50 μm (A). Box plots show median and interquartile range. Whiskers extend to 1.5x the interquartile range. B, C, D, Welch’s t-test.

### Dpy and Pio co-localise at the apical membrane of LECs and in the aECM

As Dpy and Pio interact during tubular organ morphogenesis (17–19), we studied their localisation in LECs, using Dpy::YFP together with an antibody recognising the extracellular ZP domain of Pio (17). In stationary LECs, both Dpy and Pio displayed apical punctate patterns (Fig. 5A; see also Dpy::YFP in Fig. 2C). In addition, Dpy::YFP filaments were observed, which also co-localised with Pio (*N*=49/52 cells in 5 pupae; Fig. 5A). Similar to Dpy::YFP (Fig. 1C), Pio was absent near the cell junctions (Fig. 5A). Orthogonal views showed that Pio and Dpy co-localised at the apical LEC membrane (Fig. 5A). Furthermore, Dpy localised to the aECM, where we also detected a weak and diffuse Pio signal (Fig. 5A). Imaging with the adherens junction marker Shotgun (Shg)::GFP confirmed this localisation (Suppl. Fig. S5A).

**Figure 5.**
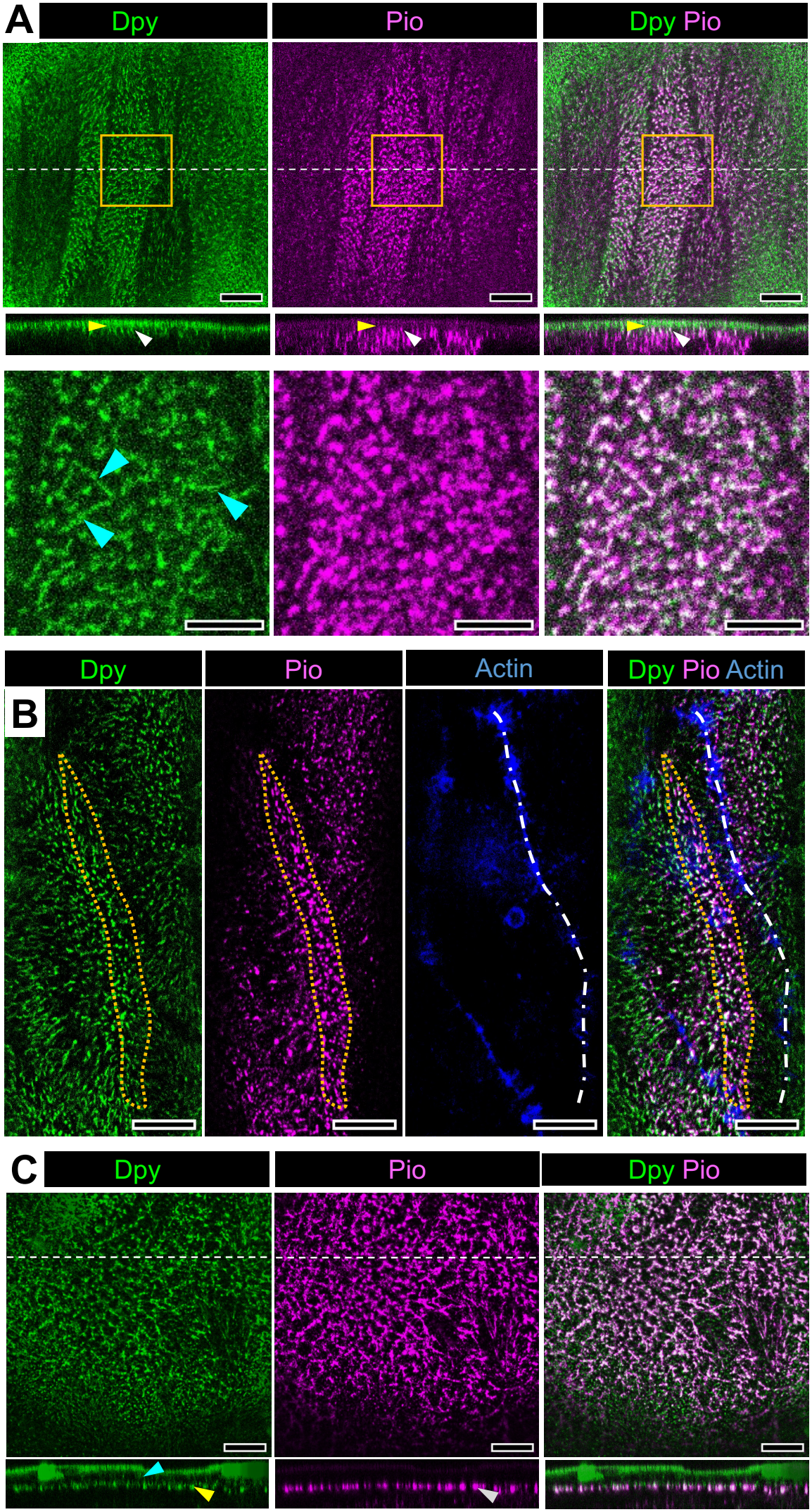
Pio co-localises with Dpy::YFP throughout abdominal morphogenesis. Anti-Pio antibody staining (magenta) in Dpy::YFP (green) pupae. **(A)** Pio co-localises with Dpy::YFP in stationary LECs (*N*=52 cells in 5 pupae). Both show apical punctate patterns (top row). However, frequently, Dpy::YFP filaments can be observed (cyan arrowheads in bottom row, which shows a magnified view of the orange box in the top row, *N* = 49/52 cells in 5 pupae). *XZ*-orthogonal views (centre row) show that the co-localisation of Dpy and Pio occurs close to the apical membrane (white arrowheads). Furthermore, Dpy localises to the aECM, which also exhibits a weaker and more diffuse Pio signal (yellow arrowheads **(B)** In early-stage migratory LECs, both Dpy::YFP and Pio exhibit a punctate pattern, with co-localisation predominantly detected in a narrow band (orange dotted outline) just behind the lamellipodium (white hatched line) (*N*=19 cells in 2 pupae). Cell outline labelled by actin staining with phalloidin (blue). **(C)** In late migratory LECs, Dpy::YFP and Pio co-localise along a fibrous network (*N*=3 pupae). The orthogonal views in the bottom row show both the remains of the early layer and the late layer of Dpy::YFP (cyan and yellow arrowhead, respectively). Pio localises with the late Dpy::YFP layer (grey arrowhead). In all images, anterior is to the left. White dashed lines indicate region of *XZ*-orthogonal views shown in A, C. Scale bars, 10 µm.

In LECs that were beginning to change shape just before they became migratory or underwent early apoptosis, highly organised Dpy::YFP filaments were visible along the cell edges, with Pio co-localising with approximately one-third of the Dpy::YFP filaments (Suppl. Fig. S5B; *N*=19/29 cells in 3 pupae). The orientation of the filaments suggests that they arose due to tension on Dpy, generated by the cell failing to detach from the aECM in time for changing their shape. Furthermore, the positioning of Pio in relation to the Dpy::YFP filament suggests that Pio co-localises with the basal part of the filaments at the apical membrane.

In the early stages of LEC migration, Pio and Dpy::YFP displayed a punctate pattern, with co-localisation primarily observed in a narrow band just behind the lamellipodium (Fig. 5B).

In late migratory LECs, Dpy::YFP and Pio co-localised along a fibrous network in the aECM (Fig. 5C; see also Dpy::YFP in Fig. 2C’).

Taken together, these findings show that Pio co-localises with Dpy at the apical membrane and in the aECM of stationary and migratory LECs, with their spatial pattern changing as cells transition from a stationary to a migratory state.

### Dpy and Pio regulate each other’s localisation and *dpy* regulates *pio* expression

The co-localisation of Dpy and Pio suggests that they might interact. To further explore this potential interaction, we studied Dpy and Pio localisation in *dpy-RNAi* and *pio-RNAi* pupae as well as in the double knock-down background (*dpy-RNAi, pio-RNAi*). We expressed the dsRNA in the P compartment and used the A compartment as internal control.

In d*py-RNAi* LECs, Dpy::YFP is strongly reduced in both the aECM and the cytoplasm in the P compartment, illustrating knock-down compared to the control (Fig. 6A, B, I). Pio is strongly reduced from the aECM and cytoplasm, compared to control pupae (Fig. 6E, F, J) and the internal control (the A compartment) (Fig. 6F). This suggests that the expression of *pio* in LECs is Dpy-dependent.

**Figure 6.**
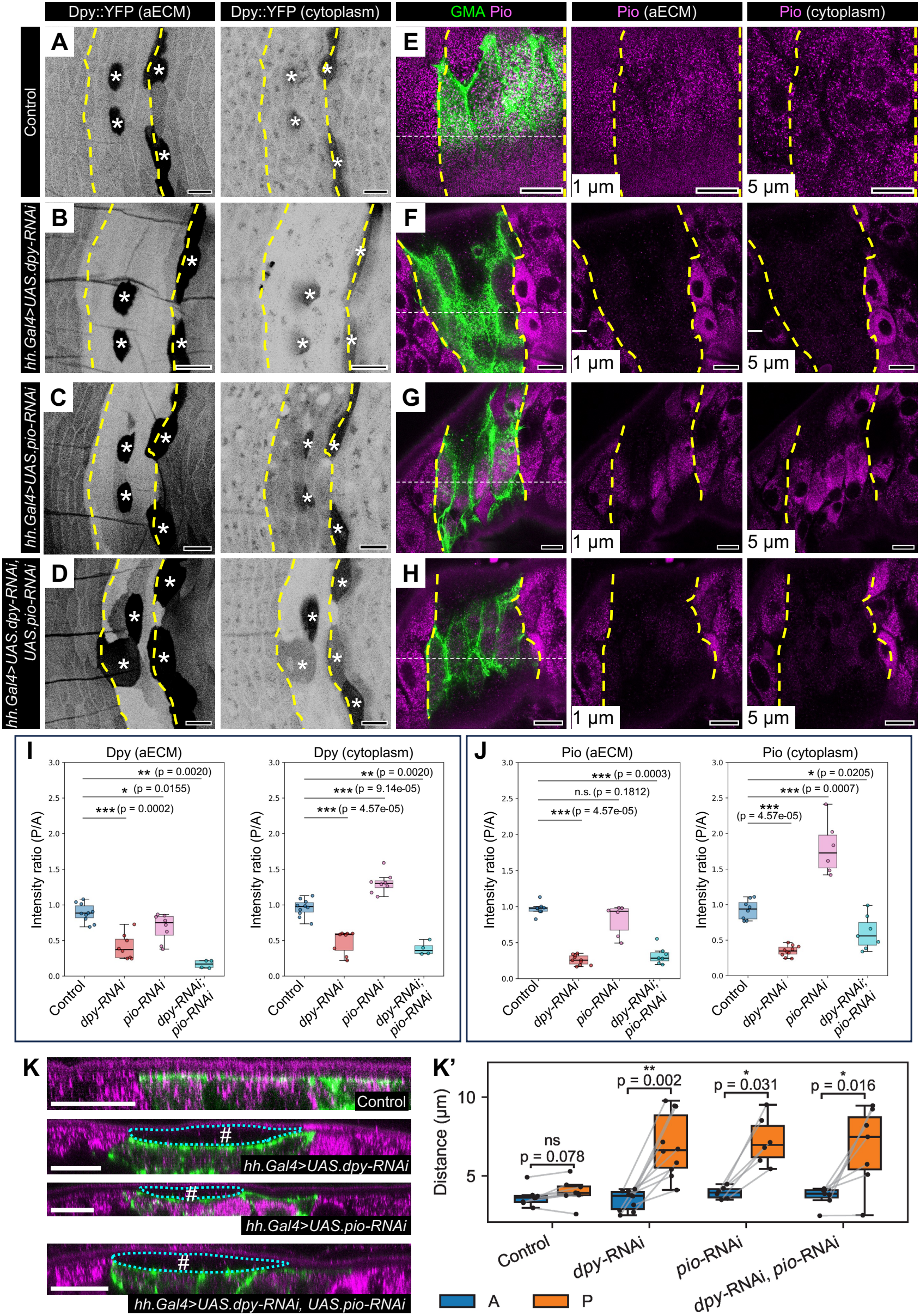
Localisation of Dpy and Pio in early, stationary *dpy* and *pio* knock-down LECs. LifeAct-Ruby and RNAi constructs were expressed in the P compartment with *hh*.*Gal4*. Yellow dashed lines indicate P compartments. Tendon cells are marked with asterisks. (A-D) Dpy::YFP localisation in aECM and cytoplasm. **(A)** Control: Dpy::YFP distribution is similar in A and P compartment LECs, in both aECM and cytoplasm (*N*=10 pupae). **(B)** *dpy-RNAi*: Dpy::YFP decreases in the aECM and cytoplasm of P compartment LECs (*N*=10 pupae). **(C)** *pio*-RNAi: Dpy::YFP is reduced in the aECM of P compartment LECs but accumulates in the cytoplasm (*N*=8 pupae). **(D)** Double knock-down (*dpy-RNAi, pio-RNAi*): Dpy::YFP decreases in both aECM and cytoplasm of P compartment LECs (*N*=4 pupae). (E-H) Anti-Pio immunofluorescence with GMA-GFP marking P-compartment LECs. **(E)** Control: Pio localises to the aECM and cytoplasm with no difference between A and P compartment LECs (*N*=8 pupae). **(F)** *dpy*-RNAi: Pio decreases in aECM and cytoplasm of P compartment LECs (*N*=10 pupae). **(G)** *pio*-RNAi: Pio is unchanged in the aECM of P compartment LECs but increases in cytoplasm (*N*=6 pupae). **(H)** Double knock-down (*dpy-RNAi, pio-RNAi*): Pio decreases in both aECM and cytoplasm of P compartment LECs (*N*=7 pupae). **(I, J)** Quantification of the ratio of relative fluorescence of Dpy::YFP and anti-Pio in A *vs*. P compartment for aECM and cytoplasm in *dpy, pio* and double knock-down pupae. **(I)** P/A intensity ratio of Dpy::YFP in aECM and cytoplasm. **(J)** P/A intensity ratio of anti-Pio in aECM *vs*. cytoplasm. **(K)** Analysis of cell detachment in *XZ*-orthogonal sections of anti-Pio immunofluorescence with GMA-GFP marking P compartment LECs. White dashed lines in E-H indicate location of *XZ*-orthogonal section. Cyan dotted lines and white hashes indicate cell detachment. Control: LECs remain attached to the aECM (*N*=7/8 pupae). In the knock-down pupae, P compartment LECs detach from the aECM (*dpy-RNAi* (*N*=7/10 pupae), *pio-RNAi* (*N*=5/6 pupae), double knock-down (*dpy-RNAi, pio-RNAi; N*=3/7 pupae)). **(K’)** Quantification of the distance between cuticle and the apical cell surface, comparing A and P compartment LECs of the experiments shown in K. Statistical significance was determined using Mann-Whitney U test (I, J) and Wilcoxon signed-rank test (K’). In all images, anterior is to the left. Scale bars, 30 μm (A-D); 20 μm (E-H, K).

In *pio* knock-down LECs, Dpy::YFP was absent from the aECM, however, the cytoplasmic Dpy::YFP signal in the P compartment was much stronger than in A compartment LECs, where *pio-RNAi* was not expressed (Fig. 6C, I). This suggests that Pio is required for transport of Dpy to the apical membrane and/or Dpy secretion into the aECM. However, unexpectedly, in P compartment LECs (*pio-RNAi*), anti-Pio antibody staining remained unchanged in the aECM and was markedly elevated in the cytoplasm compared to A compartment LECs (internal control) (Fig. 6G, J).

In the double knock-down of *dpy* and *pio*, both Dpy and Pio levels were strongly reduced in both aECM and cytoplasm (Fig. 6D, I, H, J). The phenotype resembles the *dpy-RNAi* phenotype. The cytoplasmic accumulation of Pio, which occurred when only *pio-RNAi* was expressed, disappeared in the double knock-down. This implies that Pio accumulation depends on the Dpy.

In conclusion, these data suggest that, in LECs, Dpy and Pio regulate each other.

### Knock-down of *dpy* and *pio* affects cell-aECM adhesion in stationary LECs

Interfering with *dpy* and *pio* not only disrupted LEC migration but also affected how cells adhered to the aECM (Fig. 6K). Confocal imaging of fixed stationary LECs at a small z-interval (0.19 µm) showed that control cells remained attached to the aECM (n=7, p=0.0781; Fig. 6K, K’). In contrast, *dpy-RNAi* caused cells to detach from the aECM creating a gap between aECM and LECs when expressed in the P compartment, compared to cells in the A compartment (internal control) which remained attached (n=10 pupae, p=0.00195; Fig. 6K, K’). A similar detachment was seen in *pio-RNAi* LECs (n=6 pupae; p=0.0312; Fig. 6K, K’) and in double knock-down LECs (*dpy-RNAi* and *pio-RNAi*; n=7, p=0.0156; Fig. 6K, K’)

We also observed LEC detachment in 23% of pupae expressing *pio-RNAi* in all LECs using a FLP-out cassette with *tub*.*Gal4* (*n*=13 pupae; Suppl. Fig. S6).

These results indicate that both Dpy and Pio are necessary for stationary LECs to stay attached to the aECM.

## Discussion

### Changes in LEC behaviour are accompanied by a dynamic remodelling of the aECM

In this study, we show that the LEC’s transition from stationary to migratory behaviour (33) is accompanied by a dynamic remodelling of the cells’ aECM (Fig. 7A). In stationary LECs, an apical early layer of Dpy::YFP displays a mostly punctate pattern and co-localises with Pio (Figs. 1C, D; 2A; 5A). When the LECs become mobile, they leave a footprint of their old adhesion sites behind (Figs. 1D; 2A, B), suggesting that Dpy is cleaved, releasing adhesion to allow the LECs to move. Proteolytic cleavage of Dpy as well as Pio has been observed in various morphogenetic processes. During salivary gland morphogenesis, the cleavage of the aECM by the metalloprotease ADAMTS detaches salivary gland cells, thereby promoting tube elongation during collective cell migration (43). Furthermore, Pio cleavage by protease Notopleural (Np) is involved in the formation of a luminal scaffold that is necessary for generating a uniform salivary gland tube lumen diameter (19). Cleavage of Dpy by the proteases Np and Stubble (Sb) is involved in wing development, where degradation of the aECM that surrounds the developing pupal wing is necessary for wing elongation (21), and where patterned degradation of Dpy guides the position and direction of tissue bending during wing morphogenesis (22). During tracheal development, the cleavage of Dpy by Np and Tracheal-prostatin (Tpr) is important for maintaining barrier function (44). Np-mediated ectodomain shedding of Pio releases Pio-mediated attachments between membranes and aECM (18).

**Figure 7.**
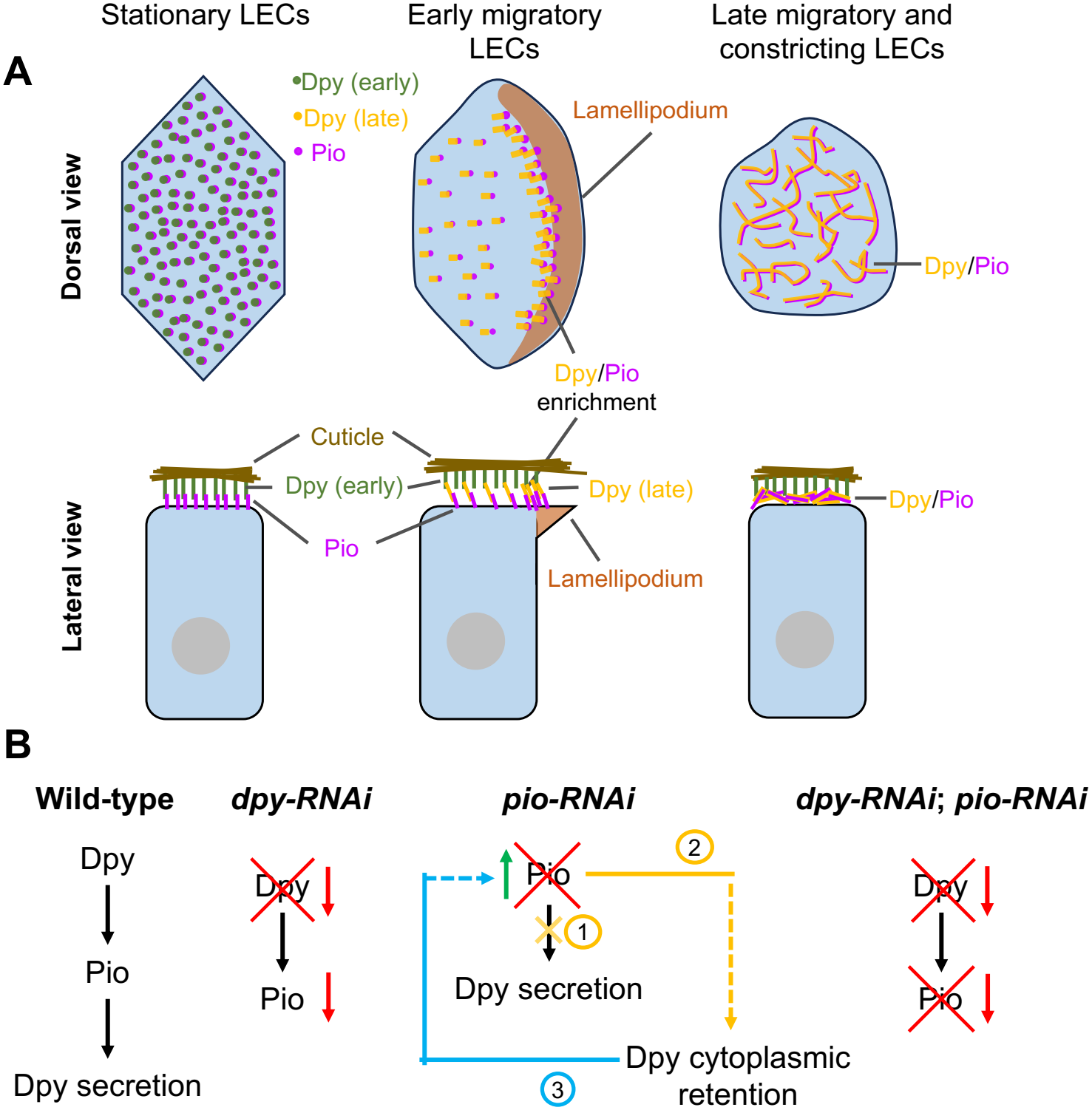
Scheme of LEC interaction with a dynamically remodelling aECM and genetic interactions of Dpy and Pio. *(Legend continued on following page)* (*Fig. 7 legend continued*) **(A)** Dorsal view: In the stationary phase, the early layer of Dpy and Pio shows a predominantly punctate pattern. During early migration, co-localisation of Dpy and Pio occurs mainly in a narrow band just behind the lamellipodium. During late migration and constriction, Dpy::YFP and Pio co-localise along a fibrous network in the aECM. Lateral view: Stationary LECs adhere to the early Dpy layer. Once LECs become mobile, they leave behind the footprints of their previous adhesion sites and secrete a new, late layer of Dpy. Gradually, this newly formed layer becomes more fibrous. **(B)** Genetic interactions between *dpy* and *pio*. Wild-type: Dpy is required for *pio* expression, and Pio is required for the apical localisation or secretion of Dpy. *dpy-RNAi*: Knock-down of *dpy* (red cross) causes a strongly reduction in *dpy* and *pio* expression (red arrows). *pio-RNAi:* Partial knock-down of *pio* may disrupt the apical localisation/secretion of Dpy (1), leading to Dpy retention in the cytoplasm (2). The remaining Pio protein may interact with Dpy in the cytoplasm (3) and accumulate there (green arrow). *dpy-RNAi, pio-RNAi*: Double knock-down of *dpy* and *pio* (red crosses) results in a strong decrease in both gene’s expression (red arrows). This phenotype is a similar to that observed in the *dpy* single knock-down.

Migrating LECs secrete a new, late layer of Dpy::YFP (Figs. 2A, B, C’; 3A’). Here, Pio colocalises with Dpy::YFP and, over time, the layer becomes more fibrous (Suppl. Movie S1; Fig. 5C). This change in morphology could reflect the deposition of new aECM by moving cells (45). It may also be associated with alterations in the mechanical properties of the aECM.

The remodelling of bECM has been extensively studied and is crucial for development and dysregulated in disease (reviewed in (46,47)). Similarly, spatio-temporal remodelling of the aECM has been observed in various developmental processes. For example, Dpy reorganizes into filament-like structures in response to mechanical stretch during indirect flight muscle development (16). During wing (20–22), tracheal (17,18), and salivary gland morphogenesis (19), the aECM undergoes dynamic turnover and eventual degradation. However, during abdominal morphogenesis it is not degradation but a change in the makeup of the aECM that takes place.

### Dpy and Pio interact genetically during abdominal morphogenesis

As Dpy and Pio colocalise throughout abdominal morphogenesis (Fig. 5A-C), we explored whether they interact. Our results suggest that *dpy* regulates the expression of *pio* (Figs. 6F, J; 7B). This result is further supported by the double knock-down of *dpy* and *pio* (Figs. 6D, I, H, J; 7B). These findings supports earlier studies, showing that Dpy and Pio physically interact during tracheal morphogenesis (18).

On the other hand, we show that apical localisation of Dpy is dependent on Pio, which appears to be required for transport of Dpy to the apical membrane and/or Dpy secretion into the aECM (Figs. 6C, I; 7B). Such a role of Pio has been described for trachea and salivary gland development (17–19). In tracheal cells, Dpy secretion depends on Tango1, which localises to specialised ER exit sites to regulate Dpy delivery from ERs to Golgi during the secretion of bulky cargo (48). In addition, post-transcriptional modification of Dpy, regulated by EOGT, is also required for Dpy secretion (49). A ZP domain-containing protein, Quasimodo (Qsm), is hypothesised to serve as a molecular chaperone, regulating the conformational maturation of Dpy protein, as well as Dpy secretion (16).

Unexpectedly, Pio accumulated in the cytoplasm of *pio-RNAi* LECs and some Pio was found at the apical membrane (Fig. 6G, J). This did not occur in the double knock-down of *dpy* and *pio* (Fig. 6D, I, H, J), indicating that Pio accumulation depends on Dpy. A possible explanation is that *pio* knock-down was partial, creating a hypomorph sufficient to produce the *pio* and *dpy* phenotypes but not entirely suppress Pio protein production. Since *pio-RNAi* retains Dpy in the cytoplasm (Fig. 6C, I), residual Pio may interact with cytoplasmic Dpy, resulting in cytoplasmic accumulation of Pio (Fig. 7B). In addition, some of the residual Pio reaches the apical membrane (Fig. 6G, J). Another possible explanation is that the anti-Pio antibody might not be specific for Pio. However, Woodward et al. (2025) showed in the salivary gland that the antibody is specific (19). A cross-reaction with Dpy is also unlikely, because the peptide sequence used to create anti-Pio does not overlap with Dpy (17), and because the staining patterns of anti-Pio and Dpy::YFP differ (Fig. 5A, B; Suppl. Fig. S5B). The used *pio-RNAi* line (VDRC_108534) has been shown to knock-down *pio* in tracheal terminal cells (48).

### A role of the aECM in anchoring stationary LECs

We have shown that the early layer of Dpy is used by the stationary LECs to anchor themselves to the aECM. Knock-down of either *dpy* or *pio* leads to cells falling off their aECM (Fig. 6K, K’; Suppl. Fig. S6). This is consistent with the primary role attributed to Dpy and Pio in mediating cell-matrix adhesion across various biological contexts, such as *Drosophila* larval development (26), epidermis-muscle attachment (14,16), wing epithelium apposition (25), as well as tracheal (17,18), salivary gland (19), wing (20–22) and posterior lobe morphogenesis (50).

Our observation of Dpy filaments (Fig. 5A) suggests that LECs apply pulling forces on their cell-aECM contacts. As LECs change shape during the transition to migratory mode, occasionally cells with aligned Dpy filaments are observed (Suppl. Fig. S5B). These cells may fail to release aECM adhesion in time, leading to force accumulation that results in the formation of Dpy filaments. Tension-induced formation of Dpy filaments has been reported in tendons linking the developing indirect flight muscles to the pupal cuticle (16).

### A role for the aECM as a substrate for LEC migration

During posterior migration, LECs secrete a new, ‘late’ layer of Dpy (Fig. 2A, C’). This layer forms beneath the footprints of the early Dpy layer and is morphologically different, appearing fibrous rather than punctate like the early layer (Fig. 2C, C’). Our results demonstrate that this aECM is needed for LEC migration. Firstly, our data show that LECs interact with the late Dpy layer. During late migration, LECs create a lamellipodia-shaped Dpy wave in front of the lamellipodial edge (Fig. 2D-G; Suppl. Movies S1, S2). This local deformation of the aECM may be due to traction forces that the LECs create at the leading edge of the lamellipodium, in line with similar findings in fibroblasts and lens corneal cells (51,52). Furthermore, LECs deform the aECM when they undergo pulsed contractions, with deformations of the late Dpy::YFP layer correlated with the periodic assembly and disassembly of actin foci in the pulsing cells (Suppl. Fig. S1; Suppl. Movie S3). This demonstrates that LECs adhere to the aECM and exert forces which deform it.

Secondly, knock-down of *dpy* and *pio* impairs LEC migration speed, migrated distance and directionality of movement (Figs. 3, 4). The decreased speed and migration distance suggest that the cells exhibit reduced traction. Deviation from the posterior migration direction is likely a consequence of the reduced migration speed and distance, as LECs begin dorsal movement in a more anterior position, thus, showing less posteriorly directed migration and more dorsal movement. As Pio knock-down affects transport of Dpy to the apical membrane and/or its secretion (Fig. 6C, I), the observed effects of *pio-RNAi* on LEC migration could be indirect via a reduction of Dpy in the aECM.

Taken together, our data provides evidence that the aECM can support cell migration. While the bECM has long been recognised as essential for cell migration in numerous contexts (53,54), such a role of the aECM as a migratory substrate has received comparatively little attention. A recent preprint showing that Dpy is involved in the migration on the aECM of cells in the *Drosophila* pupal notum (55) offers further support for this largely unexplored role of the aECM.

LECs migrate by forming crescent-shaped lamellipodia (Figs. 1A, 5B, 7) (33). In many systems, cell locomotion through lamellipodia depends on basal adhesion to the substratum at the leading edge, followed by disassembly of adhesion at the trailing edge (56). A similar mechanism may be involved in LEC migration. Future studies are required to determine how LECs interact with the aECM during their migration.

At the onset of migration, LECs show an enrichment of Dpy and Pio in a crescent-shaped region located just behind the lamellipodium (Fig. 5B). This structure may be equivalent to the actin arc, which associates with focal adhesions in the lamella, situated behind the lamellipodium, during integrin-based cell migration (57). Such a structure could be necessary for stabilising the lamellipodium during its formation as LECs begin to migrate.

### A switch from an anchoring scaffold to a substrate for cell migration

Our findings demonstrate that changes in LEC behaviour are accompanied by a dynamic remodelling of the aECM. During their stationary phase, LECs adhere to the aECM. Once they become migratory, they detach and secrete a new layer of aECM which supports their migration. These results broaden our understanding of the diverse roles the aECM plays during morphogenesis and underscore that adhesion to the aECM can be highly dynamic.

## Materials and Methods

### Fly stocks

Flies were maintained at 25ºC on standard culture medium. FlyBase (58) entries of the used transgenes are as follow:

*tub-FRT-CD2-FRT-Gal4: Rnor\CD2*^*A902*^;

*UAS*.*mCD8-GFP: Mmus/Cd8a*^*Scer\UAS*.*T:Avic/GFP*^, *UAS*.*FLP: FLP1*^*Scer\UAS*.*cDa*^;

*hs*.*FLP: FLP1*^*hs*.*PS*^;

*tub*.*Gal80ts: Scer\Gal80*^*ts*.*αTub84B*^; *hh*.*Gal4: Scer/Gal4*^*hh-Gal4*^;

*UAS*.*GMA-GFP: Moe*^*Scer\UAS*.*T:Avic\GFP-S65*^ (59);

*UAS*.*LifeAct-Ruby: Scer\ABP140*^*Scer\UAS*.*T:Disc\RFP-Ruby*^ (60);

*Dpy::YFP: dpy*^*CPTI001769*^ (61);

*Viking::GFP*: vkg^G00454^ (62);

*UAS*.*dpy-RNAi* (BSCB_31966 (63));

*UAS*.*pio-RNAi* (VDRC_107534 (64)).

### Expression of UAS-transgenes in LECs

The FLP-out technique (41) was used to overexpress UAS-transgenes in all LECs (31). Clones in the polyploid LECs, but rarely in the diploid histoblasts, were induced by heat-shocking late third instar larvae for 15 min at 37ºC. Additional expression of UAS.FLP increased the recombination events in the polyploid cells, thus increasing expression levels (31). After heat-shock, flies were kept at 25ºC for imaging.

Expression of transgenes in P compartment LECs was driven by *hh*.*Gal4*. Transgene expression was repressed using *tub*.*Gal80ts* (65) until late third instar larval stage, when larvae were shifted from the permissive (25ºC) to the restrictive temperature (29ºC) to allow Gal4-driven expression. Longer incubation at 29ºC resulted in some cases in stronger phenotypes (see results section). If the normal procedure resulted in failure of pupation due to a strong early phenotype, larvae were kept at 18ºC before heat shock or shifting to 29ºC to suppress leaky expression of the Gal4. LifeAct-Ruby (60) and GMA-GFP (59) are actin reporters that were used to label P compartment LECs, highlighting cell outlines.

Genotypes used in this study are shown in Suppl. Tab. S1. Pupae carrying clones that expressed the UAS-transgenes (UAS.X) marked with *UAS*.*mCD8-GFP* or *UAS*.*LifeAct-Ruby* had the following genotypes:

1. *y, w, hs*.*FLP; UAS*.*mCD8-GFP, tub-FRT-CD2-FRT-Gal4, UAS*.*FLP/+ or Sp* or *UAS*.*X; +* or *MRS/+* or *MRS* or *UAS*.*X*
2. *w; UAS*.*LifeAct-Ruby, Dpy::YFP/+* or *Sp* or *UAS*.*X; hh*.*Gal4, tub*.*Gal80ts/+* or *MRS* or *UAS*.*X*

Knock-down efficacy of the *dpy-RNAi* (BDSC_BL31966) line was assessed by studying protein levels using Dpy::YFP (Fig. 6C,D). Efficient *pio* knock-down by the *pio-RNAi* (VDRC_107534) line has been demonstrated in tracheal cells (48).

### 4D confocal microscopy

Pupae were prepared for microscopy as described previously (66). Pupae were staged according to (67). Sex was not determined. All pupae in the *in vivo* imaging experiments developed into pharate adults or hatched. The analysis focused on the abdominal segment 2 (A2). Imaging was done using a Leica TCS SP8 confocal laser-scanning microscope (Leica Microsystems, Mannheim, Germany), using HCPL APO CS2 40×/1.30 oil or HCPL APO CS2 63×/1.40 oil objectives at 25±2ºC. *Z*-stack step sizes were 1-2 µm. Time intervals for *in vivo* imaging were 180 or 185 s. All images and movies shown are *z*-projections of the recorded stacks. *XZ*-orthogonal views were generated in Fiji (68) using the Reslice function. In some cases, as indicated in the figure legends, z-projections of five consecutive Reslice-outputs have been used.

### Image analysis

Figures and movies were made using Microsoft PowerPoint, Adobe Illustrator, Fiji (68).

### Quantification of fluorescence intensity over time in early and late layer of Dpy::YFP

The data was aligned in time using the onset of posterior migration, which was defined as 0 hrs. The fluorescence intensity of Dpy::YFP was quantified in a region of interest (approximately 60 × 60 μm) at the *z* levels corresponding to the aECM and aECM/cell interface, respectively, using Fiji (68). Dpy::YFP intensities were measured 2 hrs prior to the onset of posterior migration and subsequently at hourly intervals until the onset of dorsal migration. The intensity at-2 h was used for normalisation. The normalised intensity was plotted against time (Fig. 2B).

*Quantification of the relationship between Dpy::YFP density and lamellipodial position* To investigate the relationship between lamellipodial position and Dpy::YFP density, regions of interests (ROI) were selected in 4D movies where the lamellipodia of several posteriorly migrating LECs were visible. ROIs were chosen such that cell migration occurred predominantly in the horizonal (x-) direction. All ROIs occupied 10 µm in the vertical (y-) direction. This ensured that they contained all but the very edges of the lamellipodia. In each ROI, lamellipodia were manually annotated as a line of 1 pixel width. 27 cells in six ROIs in two pupae were analysed. Time intervals between frames were 180 and 185 s, respectively.

(a)Kymographs (Fig. 2E): Kymographs were generated by averaging Dpy::YFP intensity values along the vertical (y-direction) for each horizonal (x-) position at each time point. In the kymographs, lamellipodial positions were displayed in pink as average position of the annotated lamellipodium at each time point.

(b)Average Dpy::YFP intensity profile relative to lamellipodial edge (Fig. 2F, G): Before our analysis, we smoothened each video frame with a Gaussian filter (σ = 3 pixels). Each annotated lamellipodium was analysed individually to estimate average Dpy::YFP intensities relative to the lamellipodium position. Specifically, for each annotated pixel, Dpy::YFP intensity was recorded along a line extending horizontally for 101 pixels along the x-axis, spanning 50 pixels on either side of the lamellipodium annotation. Each recorded line had a height of one pixel. This produced 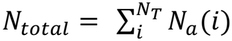 intensity profiles, where *N*_*T*_ is the number of timeframes for which the lamellipodium was annotated, and *N* _α_ (*i*) is the number of annotated pixels for the lamellipodium at time frame *i*. All recorded intensity profiles were aligned horizontally so that the lamellipodium position was at the horizontal position *x* = 0, ensuring that all recorded profiles shared identical, centred x-coordinates. To make intensity values comparable across all recorded profiles, we also subtracted from each profile the intensity value *I*(*x* = 0), the Dpy::YFP intensity at the annotated pixel of the lamellipodium position. Finally, for each of the 101 available horizontal pixel distances from the lamellipodium, all *N*_*total* intensity values were averaged. This resulted in an average profile across all annotated pixels and all time frames available for the lamellipodium. In addition to the average intensity calculated in this way, we also calculated the 50% and 90% equal-tail intervals at each pixel distance, visible as shaded areas in Fig. 2G. From the average intensity profile, we further identified peak locations using the find_peaks function from the scipy python package (69). Only the closest peak ahead of the lamellipodium was recorded. Profiles with peaks more than 5 µm away were classified as peak-less, which was confirmed by manual inspection for 6 of 27 lamellipodia. Fig. 2G shows data for all of the 21 lamellipodia with peaks.

### Quantitative analysis of cell migration in hh.Gal4 movies

To quantify the migratory behaviour of LifeAct-Ruby-labelled LECs of the P compartment, the position of the anterior boundary of the P compartment at the midline was identified at the onset of posterior migration. The position of the same boundary was then recorded at the end of posterior migration. The speed of posterior migration was calculated based on the distance between these two positions and the duration of the migration.

### Quantitative analysis of cell migration in tub.Gal4 movies

To quantify the migratory behaviour of the mCD8-GFP-labelled LECs, cells were tracked manually with the software Simi BioCell (42). Only movies of pupae were analysed that met the following criteria: (1) The time interval between frames was not longer than 3 min, (2) posterior migration had not yet started, (3) all cells were visible throughout development. All visible LECs of segment A2 were tracked. The 3D position of each LECs was recorded every 15 min. A custom-built software package (32) was then used to extract the X and Y coordinates of the centroids of the tracked cells, with positions recorded in pixels. These data were used to calculate migration speed and the direction of migration. All calculations were performed in two dimensions due to the planar character of the epithelial sheet.

(1) *Migration speed:* Velocity in pixels per frame (*V*_*pf*_) was calculated, with *V*_*pf*_ = sqrt((x_2_-x_1_)^2^+(y_2_-y_1_)^2^)/frame interval. Conversion into physical units (μm/min) followed:

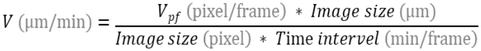

(2) *Direction of migration:* The direction of cell movement was calculated by measuring the angle formed between two successive cell positions in relation to the a–p axis of the pupa (0° corresponding to the a–p axis and 90° to the dorsal direction). Since it can be assumed that both sides of the body exhibit similar behaviour, the left and right sides were treated as equivalent, and the absolute values of the angles were used.

To visualise the mean speed and mean migration direction of all LECs across the analysed pupae, the tracking datasets for each genotype were first aligned to the onset of posterior migration. At each time point, we calculated the mean speed and mean angle along with the corresponding standard deviation. The data were then plotted over time using a dual y-axis graph, with shaded regions representing ±1 standard deviation (Suppl. Figs. S3, S4). Plots were created in Python with Matplotlib. The plots allowed us to quantify and differentiate the four phases of LEC behaviour (refer to the legends of Suppl. Figs. S3, S4 for more details).

In addition, we used the tracking data to determine the average speed and average migration direction of all tracked LECs across all time points during the identified posterior migration phase (Figs. 3C, D; 4B, C). We also calculated the posterior migration distance, normalising the data by segment size along the a-p axis (Figs. 3E, 4D).

### Quantification of Dpy::YFP and anti-Pio fluorescence intensity in A vs. P compartment

Fluorescence intensity of Dpy::YFP and anti-Pio staining was quantified by measuring mean grey values of three randomly selected ROIs per compartment (A and P) at the aECM-cell interface and 4 µm below, in the cytoplasm, avoiding nuclei (ROI sizes 50.5-422.5 µm^2^). The average of the three measurements per compartment was used to calculate the difference in fluorescence intensity between the A and P compartments, expressed as the P/A intensity ratio (Fig. 6I, J).

### Quantification of the distance between the cuticle and the apical LEC surface

To measure the distance between the cuticle and the apical surface of LECs in both control and knock-down pupae (Fig. 6K’), we manually annotated both regions in XZ sections of fixed LECs stained with anti-Pio. RNAi constructs were expressed in the P compartment using *hh*.*Gal4*. LECs of the A compartment, which did not express the RNAi construct, served as internal control. For each experimental condition, we annotated A and P compartments separately by drawing two 1-pixel-wide lines in Fiji (68): one along the cuticle and one along the apical surface of the tissue.

For each pair of annotated lines, we extracted distances as follows. First, a centreline was constructed between the two annotated lines. Specifically, each line annotation was parameterized by its normalised line length, which we will refer to as arclength, such that each arclength value between zero and one corresponded to one point on each annotated line. Then, for 300 evenly spaced values of the arclength between zero and one, the corresponding points on the two annotated lines were extracted, and their midpoint determined. Thereby, a centreline was drawn, parameterized by the arclength. Then, we fitted a straight line to this centreline, using least square optimization. We refer to this straight line as the tilt-corrected centreline. Fitting a quadratic instead of a straight line to the centreline did not change the results.

To extract distance measurements,100 evenly spaced points along the arclength of the tilt-corrected centreline were sampled. At each point, distances between the two lines were measured along a line going through the point and directed perpendicular to the tilt-corrected centreline. The average value of these 100 distances was taken as the average distance of the given annotation.

For each experimental conditions, we used a Wilcoxon signed-rank to test whether differences between A and P annotations were non-zero.

### Quantitative analysis of the similarity of aECM deformation and actin dynamics

DEFORM-Net (40) was used to quantify the displacement vector fields for both Dpy::YFP and LifeAct-Ruby fluorescence intensity time-series to assess the similarity between aECM deformation and actin dynamics of LECs that undergo pulsed contractions (Suppl. Fig. S1; Suppl. Movie S3). Each fluorescence channel of the timelapse videos was independently evaluated with a pre-trained DEFORM-Net model to generate two time series of displacement vector fields. These displacement fields were visualised as 2D vector plots, with the arrow magnitude scaled to eight times the true displacement. The vector plots corresponding to the aECM and actin were overlaid in green and orange colour, respectively, over the corresponding reference frames of the actin fluorescence intensity (grayscale).

In addition, we evaluated the dot product between the two displacement fields to quantify vector similarity across the image. The dot product is defined as:

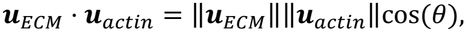

where *θ* is the angle between the vectors. A large positive value indicates aligned vectors with high magnitudes, a large negative value indicates opposite orientations, and a small value reflects low magnitudes and/or weak similarity. The dot product value was displayed over the corresponding reference frames of the actin fluorescence intensity (grayscale) as a multiplicative overlay, with red signifying a high positive value and blue signifying a high negative value in the range of [-0.9, 0.9] µm.

### Quantification of width of P compartment LECs

Width of P compartment LECs was measured using Fiji (68). A straight line was drawn along the short axis of each cell, perpendicular to its long axis, at the level of the apical cell outlines. The length of this line was measured. Statistical significance between groups was assessed using Mann-Whitney U test (Suppl. Fig. S2).

### Immunofluorescence of the pupal abdominal epidermis

Immunohistochemistry followed standard procedures (70). Pupae were staged after (67), bisected into dorsal and ventral halves, and fixed in 4% PFA (Thermo Fisher Scientific, 28908) for 20 min at room temperature (RT). Tissues were permeabilised in PBST (0.1% Triton X-100 (Sigma, X100) in PBS) for 15 min, then blocked in 1% BSA (Sigma, A2153) in PBST for 30 min at RT. Primary antibodies were incubated overnight at 4ºC, followed by 1 h secondary antibody incubation at RT with Alexa Fluor™ 635 Phalloidin (Thermo Fisher Scientific, A34054). DAPI (1 μg/mL; Thermo Fisher Scientific, D1306) was applied for 5 min at RT before mounting with VECTASHIELD® HardSet™ Antifade (H-1400-10).

Used antibodies and dilutions: Rabbit anti-Pio antibody 1:200 (17); goat Anti-Rabbit IgG H&L (Alexa Fluor® 568; Abcam, ab175471) 1:1000.

The tissue was imaged using a Zeiss LSM 800 confocal microscope (Zeiss, Jena, Germany) with a Plan Apochromat 63x/1.4 oil DIC objective. *Z*-stack imaging was applied, with various *Z*-range and *Z*-intervals (0.19-0.5 µm). Maximum-intensity projection and orthogonal *Z* analysis were applied using Fiji (68). *XZ*-orthogonal views were generated using the Reslice function. In some cases, orthogonal views were created by *z*-projecting five consecutive Reslice-images (indicated in the corresponding figure legends). Data was saved as TIFF files.

### Statistics

No analysable samples were excluded from the analysis. All statistical analyses and data visualisation were performed using Python (v3.13.0) with the following libraries: SciPy (v1.15.3) for statistical testing, NumPy (v2.1.3) and Pandas (v2.2.3) for data handling, and Matplotlib (v3.10.0) for figure generation. Quantitative data were visualised as box plots displaying the median, interquartile range, and whiskers extending to 1.5x the interquartile range. Individual data points were overlaid on each box plot.

For datasets where each group contained at least five observations, normality was evaluated using the Shapiro-Wilk test. When all groups met the normality criterion (p > 0.05), Welch’s t-test (two-sided, assuming unequal variance) was employed. If any group failed the normality test, the Mann-Whitney U test (two-sided) was used instead. In the quantification of Dpy::YFP and anti-Pio fluorescence, some groups included fewer than five observations, and the small sample sizes prevented a reliable normality assessment; here, the Mann-Whitney U test was applied. The same test was consistently applied across all comparisons. For the analysis of cell detachment (Fig. 6K’), a Wilcoxon signed-rank test was used.

Significance thresholds were defined as p<0.001 (***), p<0.01 (**), p<0.05 (*), and p≥0.05 (not significant, n.s.). Sample sizes for each experiment are indicated in the figure legends as ‘*N* = phenotype number/total number’.

## Acknowledgements

We thank Barry Denholm, Stefan Harmansa and Jens Januschke for comments on the manuscript, Markus Affolter for the gift of anti-Pio antibody, Barry Thompson and Barry Denholm for flies. Stocks obtained from the Bloomington Drosophila Stock Center (NIH P40OD018537) and the Vienna Drosophila Resource Center (VDRC, www.shop.vbc.ac.at) were used in this study. We used FlyBase (release FB2026 02) to find information on function, gene products, stocks, alleles and insertions, interactions. J.M. acknowledges funding from the China Scholarship Council (CSC).

## Author contributions

J.M. and M.B. conducted experiments, J.M.,P.W., J.K. and M.B. analysed data, J.M.,P.W., J.K. and M.B. interpreted data, M.B. supervised the research, J.M. and M.B. designed the research, J.M. and M.B. wrote the manuscript. All authors commented on the manuscript.

