## Supplementary Information for "A dynamic role for the apical extracellular matrix in cell migration during *Drosophila* abdominal morphogenesis"

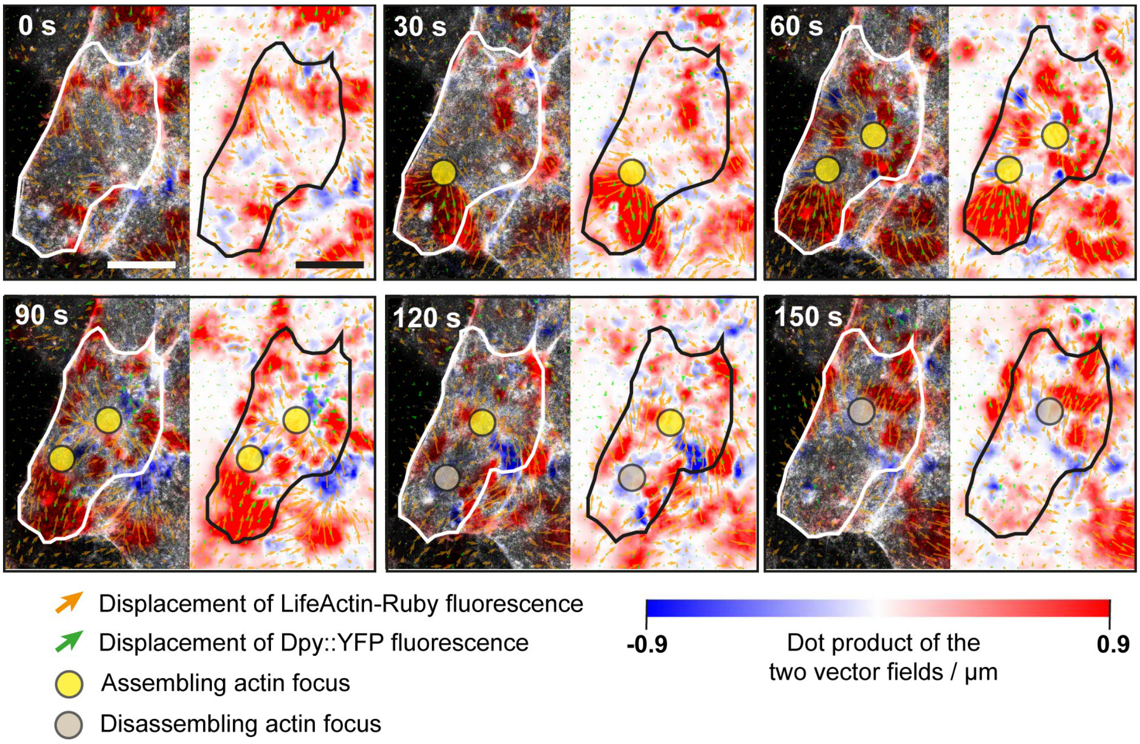


**Supplementary Figure S1. LECs that undergo pulsed contractions deform the aECM.** Six frames from a time-lapse movie showing a LEC undergoing pulsed contractions. LEC expresses Dpy::YFP and LifeAct-Ruby. Left panels: LifeAct-Ruby channel is overlaid with the DEFORM-NET displacement vector fields, with orange arrows representing LifeAct-Ruby fluorescence and green arrows representing Dpy::YFP fluorescence. The blue-white-red LUT indicates the dot product between the two vector fields. For illustration, two assembling actin foci are circled in yellow. Once they disassemble, they are marked with grey circles. Right panels: Same information is shown as in the left panels, except that the LifeAct-Ruby image is replaced by the outline of the cell. Near assembling and disassembling foci, the dot product often displays high-magnitude, aligned vectors (red), demonstrating that deformations of the late Dpy::YFP layer correlate with the periodic assembly and disassembly of actin foci in the pulsing cell (*N*=11 cells in 7 pupae). Anterior is to the left. Scale bars, 20 µm.

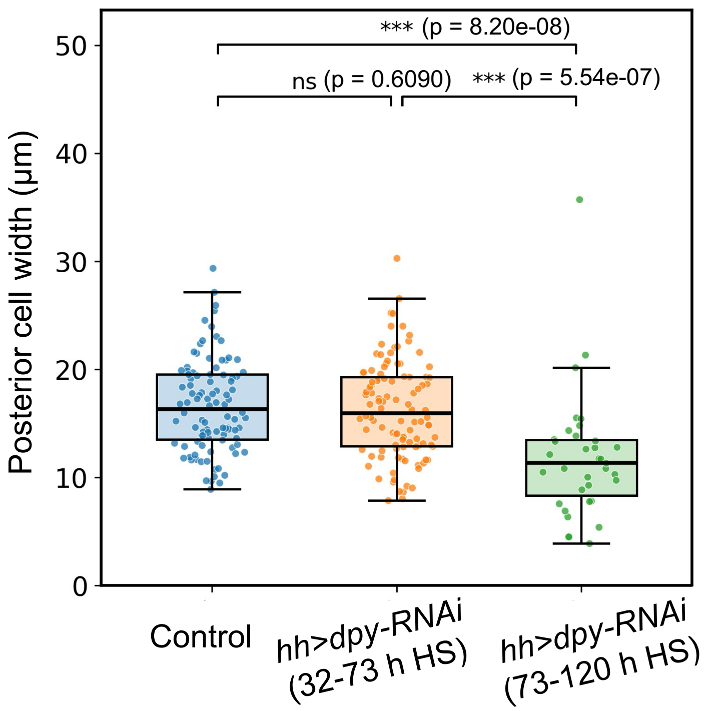


**Supplementary Figure S2. Quantification of cell width in control and *dpy-RNAi* pupae.** Strong knock-down of *dpy* results in narrower cell morphology. The strength of the phenotype corresponds to the length of time the RNAi was expressed before imaging (i.e. the length of incubation at 29ºC; 32-73 h [weaker phenotype]; 73-120 h [stronger phenotype]). Circles represent individual LECs measured in control (98 cells in 5 pupae) and knock-down pupae (106 cells in 5 pupae with weak phenotypes; 35 cells in 3 pupae with strong phenotype). Mann-Whitney U test.

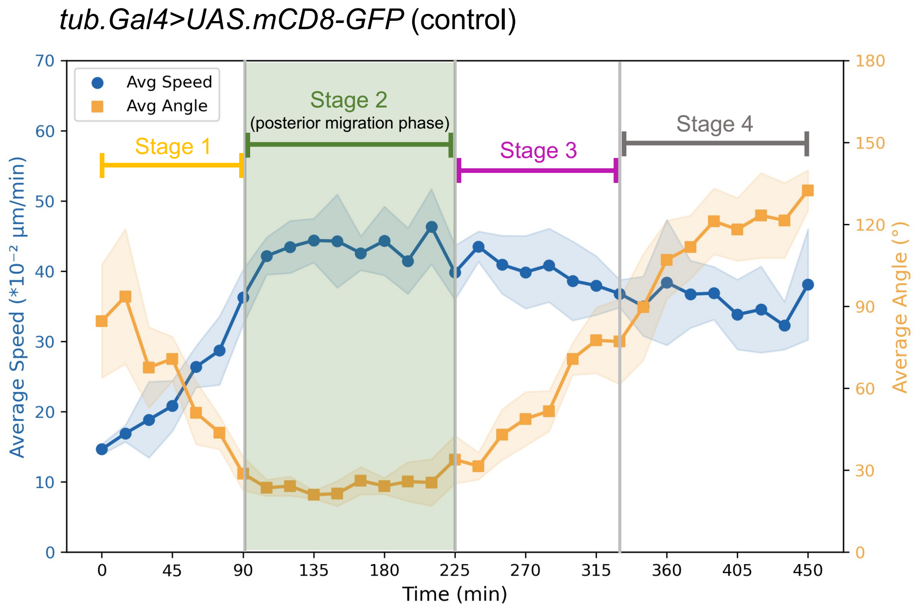


**Supplementary Figure S3. Quantitative analysis of LEC movement during abdominal morphogenesis.** LECs expressing mCD8-GFP via a tub.Gal4 FLP-out cassette were manually tracked using SIMI Biocell (42). A dual y-axis graph presents the average speed and direction (angle) of LEC movement over time, with shaded areas indicating ±1 standard deviation (*N*=6 pupae). 90° corresponds to dorsal movement towards the dorsal midline, while 0° reflects posterior movement. The plot allowed us to quantify and differentiate the four phases of LEC behaviour: *Stage 1 (0-90 min):* During the initial stages of abdominal morphogenesis, LECs transition from being stationary to becoming migratory. While some cells undergo apoptosis, the surviving cells move to occupy the spaces left behind by those that have died. Movement during this stage is limited, randomly oriented, and marked by increasing speed with variable directionality. *Stage 2 (90-225 min):* LECs migrate posteriorly (directionality ≤30°) with high and stable speeds (around 0.4 µm/min). *Stage 3 (225-345 min):* LECs change direction and migrate dorsally. At the same time, they begin to constrict, which contributes to the dorsal movement. Movement during this stage is characterised by a moderate decrease in speed and an increase in the directionality (from 30° to 90°). *Stage 4 (345-450 min):* All remaining cells constrict and delaminate, resulting in directionality values ≥ 90° and more variable movement speeds.

**A** *tub.Gal4>UAS.mCD8-GFP; UAS.dpy-RNAi*

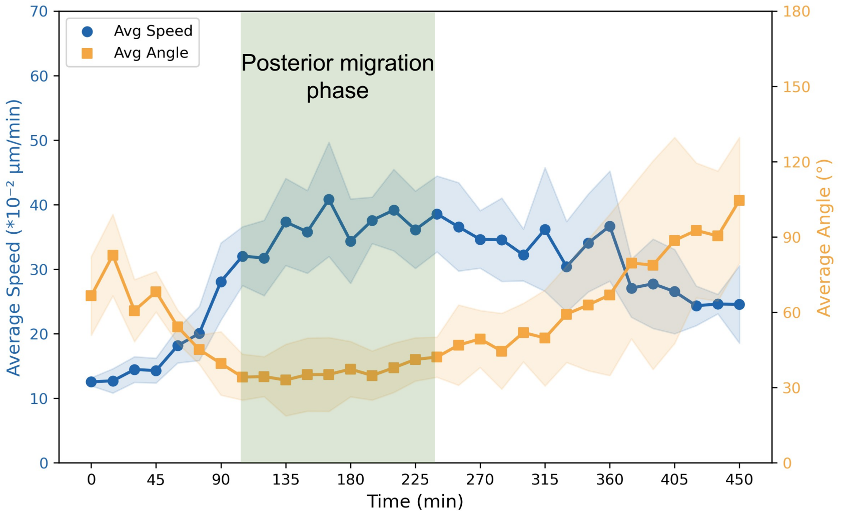


**B** *tub.Gal4>UAS.mCD8-GFP; UAS. pio-RNAi*

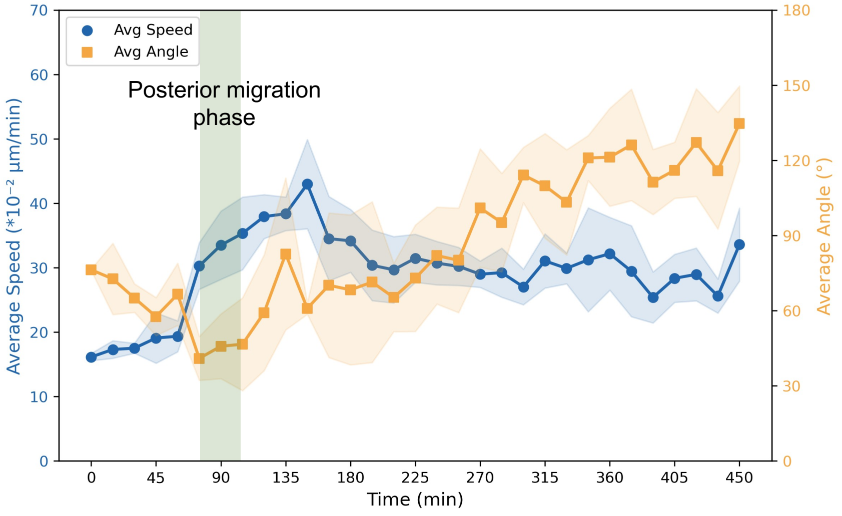


**Supplementary Figure S4. Knock-down of Dpy and Pio impairs posterior LEC migration.** LECs expressing mCD8-GFP and *dpy-RNAi* (A) or *pio-RNAi* (B) using a *tub.Gal4* FLP-out cassette were manually tracked using SIMI Biocell (42). A dual y-axis graph presents the average speed and direction (angle) of LEC movement over time, with shaded areas indicating ±1 standard deviation. 90° corresponds to dorsal movement towards the dorsal midline, while 0° reflects posterior movement. **(A)** *dpy-RNAi* (*N*=7 pupae): Compared to wild-type pupae (Suppl. Fig. S3), LECs move slower in their posterior migration phase (0.3 - 0.4 µm/min compared to ~0.4 µm/min in wild-type), and less in posterior direction (30°- 45° compared to ≤30° in wild-type). This is accompanied by greater variability, as indicated by a larger standard deviation. As in wild-type, the posterior migration phase (105–240 min) was defined as the period during which cell speed was high and directionality remained relatively constant. **(B)** *pio-RNAi* (*N*=5 pupae): Compared to wild-type pupae (Suppl. Fig. S3), the migration phase is very short (75–105 min). The phase was defined as the period during which directionality remained relatively constant and was oriented approximately posteriorly (~45°). During posterior migration, LECs exhibit slower migration speeds (~0.35 µm/min) and higher directionality (~45°). This is accompanied by greater variability, as indicated by a larger standard deviation. Thus, LECs failed to undergo prolonged posterior migration and transitioned to dorsal movement earlier than in wild-type pupae.

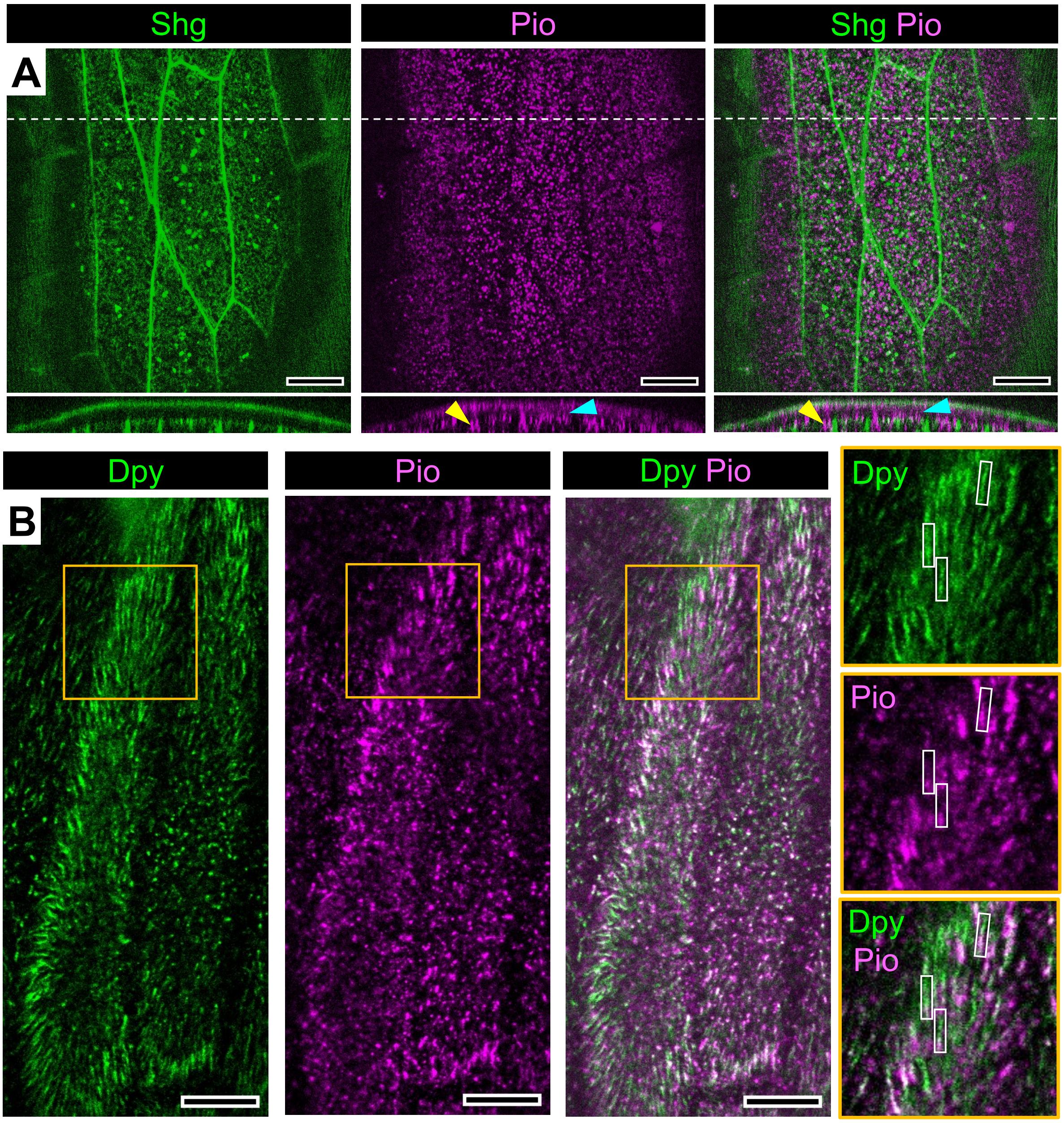


**Supplementary Figure S5. Pio localisation in stationary LECs. (A)** Anti-Pio immunofluorescence with Shg::GFP labelling adherens junctions. Shg::GFP acts as a marker for the apical membrane. Pio localises to the aECM (cyan arrowhead) and the apical cell membrane (yellow arrowhead) (*N*=43 cells in 2 pupae). Hatched white line indicates location of *X/Z* orthogonal section shown. **(B)** Anti-Pio antibody staining (magenta) in Dpy::YFP (green) pupae. In an early LEC, just as it starts to change shape before migration, Dpy::YFP filaments are visible around the edges of the cells, and anti-Pio co-localises with around one thirds of these filaments (rectangles; *N*=19/29 cells in 3 pupae). Orange box indicates zoomed in areas, which are shown on the right. In all images, anterior is to the left. Scale bars, 10µm.

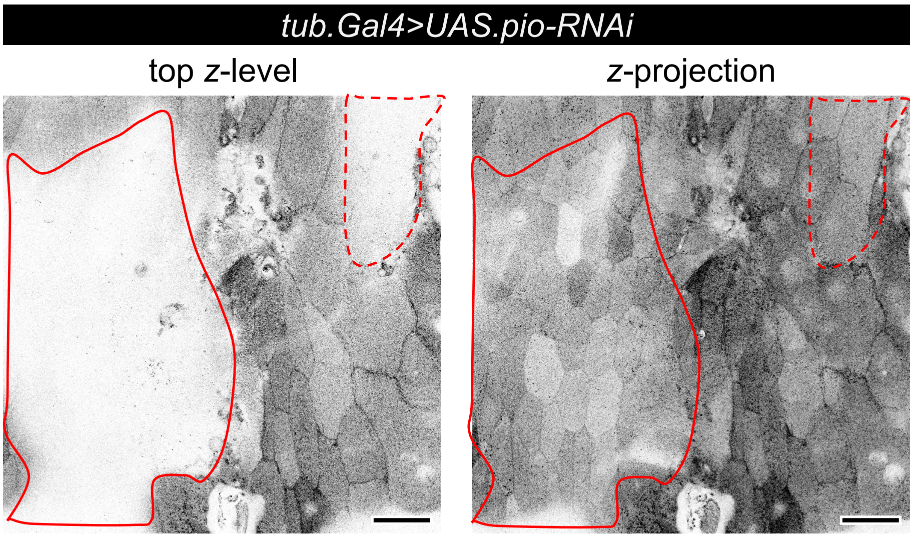


**Supplementary Figure S6. LEC detachment in *pio-RNAi* LECs.** *pio-RNAi* expressed in all LECs using a FLP-out cassette with *tub.Gal4*, and cells are labelled with mCD8-GFP. In some pupae, part of the tissue detaches from its aECM, creating a pit where the cells sink deeper into the pupae (*N*=3/13 pupae). These regions are outlined with red hatched lines. The sink is visible at the top z-level of the recorded z-stack (left), while the z-projection displays the entire tissue (right). Anterior is to the left. Scale bars, 50 µm.

**Supplementary Table S1. List of genotypes used in this study.**

| Genotype | Figure/Movie |
|---|---|
| *w; Dpy::YFP/UAS.LifeAct-Ruby; hh.Gal4, tub.Gal80ts /+* | Figs.1B,D; 2A,D; 3A; 6A; Suppl. Fig. S1, S2; Suppl. Movie S1, S2, S3 |
| *w; Dpy::YFP/Dpy::YFP, UAS.LifeAct-Ruby; hh.Gal4, tub.Gal80ts /+* | Fig.1C |
| *w; Viking::GFP/UAS.LifeAct-Ruby; hh.Gal4, tub.Gal80ts /+* | Fig.1B’ |
| *w; Dpy::YFP/Dpy::YFP; +/+* | Figs.2C; 5; Suppl. Fig. S5B |
| *w; Dpy::YFP/UAS.LifeAct-Ruby; hh.Gal4, tub.Gal80ts/UAS.dpy-RNAi* | Fig.3A’, A”; 6B; Suppl. Fig. S2; Suppl. Movie S4, S5 |
| *y, w, hs.FLP; UAS.mCD8-GFP, tub-FRT-CD2-FRT-Gal4, UAS.FLP/+; +/+* | Figs.3B; 4A; Suppl. Fig. S3; Suppl. Movie S6 |
| *y, w, hs.FLP; UAS.mCD8-GFP, tub-FRT-CD2-FRT-Gal4, UAS.FLP/Sp; UAS.dpy-RNAi/+* | Fig.3B’; Suppl. Fig. S4A; Suppl. Movie S7 |
| *y, w, hs.FLP; UAS.mCD8-GFP, tub-FRT-CD2-FRT-Gal4, UAS.FLP/ UAS.pio-RNAi; MRS/+* | Fig.4A’, Suppl. Figs. S4B, S6; Suppl. Movie S8; |
| *w; Dpy::YFP, UAS.LifeAct-Ruby/UAS.pio-RNAi; hh.Gal4, tub.Gal80ts/MRS* | Fig.6C |
| *w; Dpy::YFP, UAS.LifeAct-Ruby/UAS.pio-RNAi; hh.Gal4, tub.Gal80ts/UAS.dpy-RNAi* | Fig.6D |
| *w; UAS.GMA-GFP/+; hh.Gal4, tub.Gal80ts /+* | Fig.6E, K |
| *w; UAS.GMA-GFP/Sp; hh.Gal4, tub.Gal80ts / UAS.dpy-RNAi* | Fig.6F, K |
| *w; UAS.GMA-GFP/ UAS.pio-RNAi; hh.Gal4, tub.Gal80ts/MRS* | Fig.6G, K |
| *w; UAS.GMA-GFP/ UAS.pio-RNAi; hh.Gal4, tub.Gal80ts/UAS.dpy-RNAi* | Fig.6H, K |
| *w; Shg::GFP/ Shg::GFP* | Suppl. Fig. S5A |

### Supplementary Movies

**Supplementary Movie S1. Dpy dynamics and LEC migration during *Drosophila* abdominal morphogenesis.** Time-lapse movie showing Dpy::YFP dynamics (left) alongside LifeAct-Ruby marking P compartment LECs (right). LECs show posterior migration followed by dorsal migration and apical constriction (beginning around 6:50 hrs, mostly beyond the field of view). While a cell undergoes apoptosis (asterisk in the LifeAct-Ruby channel), the early Dpy::YFP pattern is left behind as a footprint (yellow outline in Dpy::YFP channel). Similarly, when cells begin their posterior migration, the early Dpy layer is also left behind and photobleaches over time. From around 3 hrs, the deposition of the late Dpy::YFP layer begins. The more basal late layer is visible as there is a subtle imaging drift in basal direction during the recording. The fibrous network increases in complexity over time. Crescent-shaped deformations of the Dpy::YFP network appear from around 4:42 hrs (focus on yellow box to observe Dpy::YFP waves). The interactions with the aECM cease once the histoblasts start to appear and abdominal closure takes place (from 7:03 hrs). Anterior is to the left. 30 frame per second (fps). See also Figs. 1D, D'; 2A.

**Supplementary Movie S2. During posterior LEC migration, crescent-shaped waves of high fluorescence intensity move across the late Dpy::YFP layer.** These waves travel just in front of the lamellipodia (Fig. 2D-G'). (**A**) Time-lapse movie showing Dpy::YFP (top) and LifeAct-Ruby marking P compartment LECs (bottom). Crescent-shaped waves move from anterior (left) to posterior (right). (**B**) Data for the quantification of the relationship of Dpy::YFP density and lamellipodial position (Fig. 2 F, G, G'). Time-lapse movie showing the Dpy::YFP signal, smoothened with a Gaussian filter ($\sigma$ = 3 pixels) (bottom panel). Top panel showing the smoothened Dpy::YFP signal with the manually segmented lamellipodium indicated with a cyan line. 20 fps. In both movies, anterior is to the left. Scale bars, 10 µm.

**Supplementary Movie S3. LECs which undergo pulsed contractions deform the aECM.** Time-lapse movie showing a LEC that undergoes pulsed contractions. Top left: LifeAct-Ruby labelling actin dynamics. Top right: Dpy::YFP labelling aECM deformation. Bottom right: DEFORM-NET displacement vector fields are displayed using orange arrows for LifeAct-Ruby fluorescence and green arrows for Dpy::YFP fluorescence. In addition, the blue-white-red LUT represents the dot product between these two vector fields (for LUT, see Suppl. Fig. S1). Bottom left: Overlay of LifeAct-Ruby channel, DEFORM-NET displacement vector fields, and the dot product between the vector fields. Near assembling and disassembling foci (rhythmical fluorescence accumulations in the LifeAct-Ruby channel), the dot product often shows high-magnitude, aligned vectors (red), demonstrating that deformations of the late Dpy::YFP layer correlate with the periodic assembly and disassembly of actin foci in the pulsing cell. Anterior is to the left. Scale bar, 20 µm. See also Suppl. Fig. S1.

**Supplementary Movie S4. Dpy deposition is independent of the early layer of Dpy.** Time-lapse movie of segment A2 P compartment LECs and adjoining A compartment LECs (yellow lines indicate P compartment). All LECs express Dpy::YFP; P compartment cells also express LifeAct-Ruby and *dpy-RNAi*. In this weak *dpy-RNAi* pupa, Dpy::YFP is reduced above the P compartment LECs. Posterior LEC migration takes place. The migrating A compartment LECs deposit a late Dpy::YFP layer in the area that has previously been occupied by the P compartment. Anterior is to the left.

**Supplementary Movie S5. Strong *dpy* knock-down impairs LEC migration.** Time-lapse movie of segment A2 P compartment LECs and adjoining A compartment LECs (yellow lines indicate P compartment). All LECs express Dpy::YFP; P compartment cells also express LifeAct-Ruby and *dpy-RNAi*. In this strong *dpy-RNAi* pupa, Dpy::YFP is strongly reduced above the P compartment LECs. P compartment LECs undergo little posterior movement and show a narrow morphology. Abdominal closure takes place, nonetheless. As LECs are not migrating, the area above the P compartment remains with little Dpy::YFP. Anterior is to the left.

**Supplementary Movie S6. Wild type LEC migration.** All LECs are labelled by mCD8-GFP driven by *tub.Gal4*. LECs undergo early apoptosis (asterisks) and stationary-to-migratory transition, followed by posterior migration. Around 6:09 hrs, cells change direction and move dorsally, followed by apical constriction and replacement by histoblasts. Anterior is to the left. Scale bar, 20 μm.

**Supplementary Movie S7. Knock-down *of dpy* in all LECs interferes with posterior migration.** LECs expressing *dpy*-RNAi and *mCD8-GFP* under *tub.Gal4* in all LECs. LECs initiate posterior migration (around 4:51 hrs), however, they change to dorsal migration soon after becoming mobile (around 6:39 hrs), resulting in a shorter posterior migration phase. LECs then undergo apical constriction and replacement by histoblasts. Anterior is to the left. Scale bar, 20 μm.

**Supplementary Movie S8. Knock-down of *pio* impaired posterior migration interferes with posterior migration.** LECs expressing *pio*-RNAi and *mCD8-GFP* under *tub.Gal4* in all LECs. LECs initiate posterior migration (around 1:00 hrs), however, they change to dorsal migration very soon after becoming mobile (around 3:03 hrs), resulting in a very short posterior migration phase. LECs begin apical constriction early and replacement by histoblasts occurs. Anterior is to the left. Scale bar, 20 μm.
